# Bumble Bee Abundance and Diversity Increase with Intensity of Tallgrass Prairie Restoration Intervention

**DOI:** 10.64898/2026.03.24.713996

**Authors:** Jade M. Kochanski, Stephanie L. McFarlane, Ellen I. Damschen, Claudio Gratton

## Abstract

**Introduction:** Human land-use intensification and the resulting habitat loss are primary drivers of insect pollinator declines. Habitat restoration offers a promising approach to counteract these declines, yet landscape-level evaluations of bee responses to restoration and management remain limited. We conducted a two-year, landscape-scale study in Wisconsin, USA, to assess how different intensities of tallgrass prairie restoration and management affect bumble bees (*Bombus* spp.).

**Objectives:** This study aimed to determine whether (1) bumble bee abundance and diversity increase with assisted restoration, and (2) outcomes differ between low-(seeded only) and moderate-intensity (seeded and managed with prescribed fire) interventions.

**Methods:** Using catch-and-release surveys, we measured bumble bee abundance and diversity at 32 sites representing a gradient in restoration intervention: no intervention (unassisted recovery), low intervention, and moderate intervention.

**Results:** Bumble bee abundance and diversity were higher at assisted restoration sites (low and moderate intervention) than at unassisted sites. Although both tended to be greater at moderate than low intervention intensities, these differences were not statistically significant. Bumble bee community composition also differed across intervention intensity, driven by shifts in dominant species (e.g., *B. impatiens* and *B. griseocollis*). Rarer taxa, including endangered and vulnerable species, occurred only at assisted restoration sites, with the largest populations at moderate intervention sites. Across all sites, bumble bee responses were strongly and positively associated with floral abundance, but not with semi-natural habitat in the surrounding landscape.

**Conclusion:** Our findings demonstrate that assisted grassland restoration can effectively increase bumble bee abundance and diversity, supporting its value as a conservation practice for pollinators.

Implications for Practice: (1) Grassland restorations targeting plant communities can successfully support nontarget pollinators across a range of management intensities and landscape contexts. Adding seeds of pollinator-preferred plants could improve restorations with low floral abundance and diversity. (2) Management of existing restorations is important to maintain abundant floral resources and diverse pollinator communities. Because sites varied widely in prescribed fire use, our findings likely represent a conservative estimate of its benefits, and higher intervention intensity (e.g., repeated seeding, regular fire, mechanical or chemical shrub and invasive plants control) may further enhance outcomes for bumble bees.

## Introduction

Human land-use intensification and the resulting habitat loss are primary drivers for biodiversity declines (e.g., Newbold et al. 2015; IPBES 2019). These impacts have been particularly extensive for insect pollinators, many of which are declining in population size and geographic range (Potts et al. 2010; Cameron et al. 2011; Goulson et al. 2015). For example, the U.S. federally endangered Rusty-Patched bumble bee (*Bombus affinis*) is found in only 13% of its historic U.S. range (Colla and Packer 2008). These declines are especially concerning since insect pollinators are essential components of nearly every terrestrial ecosystem. Moreover, wild, unmanaged bees are of particular importance because they pollinate 35% of crop species and 85% of all other flowering plants (Ollerton et al. 2011). Since declines in *B. affinis* and other bee species have been attributed in part to habitat loss and fragmentation (Cameron and Sadd 2020), there are growing efforts to restore native habitats to counteract bee population declines. Yet, research on how landscape-scale habitat restoration and subsequent management affect wild bee populations remains limited.

A promising approach to addressing habitat loss is the restoration of native ecosystems. In the Midwestern U.S., tallgrass prairie once spanned approximately 162 million acres of the central U.S. but is now reduced to just 3-4% of its original range (Samson and Knopf 1994). This ecosystem supports a high diversity of plants and animals, with more than 150 flowering plant species documented in restored tallgrass prairies in the Upper Midwestern U.S. (McFarlane et al. 2023). Restoring these habitats can provide essential resources for wild bees, which rely on pollen and nectar as their key food sources and could bolster their populations. Studies in the Upper Midwest U.S. associate increases in bee abundance and diversity with prairie restoration. For example, Griffin et al. (2017) observed a ~30% increase in both abundance and diversity of wild bees in tallgrass prairie restorations when compared to agricultural fields. In contrast, Tonietto et al. (2017) found little difference in bee abundance and diversity between restored prairie and former agricultural fields. Both Griffin et al. (2017) and Tonietto et al. (2017) found that bee community composition was distinct in each of these habitats which suggests species-specific responses to restoration and that additional factors also likely influence bee communities.

Habitat restoration increases bee abundance and diversity by altering resource availability, primarily by providing flowers and nesting habitat across space and time (Blaauw and Isaacs 2014; Harmon-Threatt and Chin 2016; Williams et al. 2024). Leaving land fallow, or undergoing no restoration intervention, provides opportunities for passive regeneration of floral and nesting resources for bees. Work concurrent with ours found that while the number of native plant species did not significantly differ across no intervention and active intervention sites, the quality of the plant community increased with active seeding (McFarlane et al. 2023). Thus, active seeding may be an opportunity to improve habitat for bees across space and time. Seed mixes with higher floral diversity or pollinator-preferred species likely support more abundant and diverse bee communities than no intervention (Harmon-Threatt and Hendrix 2015; Williams et al. 2015; Havens and Vitt 2016). Bee communities have long foraging seasons; some species emerge in early spring (e.g., March, April) and others remain active until the first hard frost (e.g., October). Landscapes that offer complementary and continuous foraging resources spanning early spring into the fall support more diverse bee communities (Williams et al. 2015; Mallinger et al. 2016). Further, diverse floral resources are vital for adequate nutrition (Vaudo et al. 2014, 2015, 2016) to support growth and development, life stage dependent nutritional requirements, and to self-medicate (Adler et al. 2020; Fowler et al. 2022; Malfi et al. 2023). Finally, restoring large tracts of land can enhance nesting opportunities, decrease competition for nest sites, and provide refugia from pesticide exposure in highly modified landscapes like those of the Upper Midwest (Rundlöf et al. 2022; Williams et al. 2024). Importantly, these food and nesting resources for bees are all impacted by management intensity.

Management following seeding is an important factor often overlooked in studies examining habitat restoration impacts on bees. Intentional controlled burning (i.e., prescribed fire) is the most prevalent form of grassland management and has been fundamental to maintaining prairie biodiversity for millennia (Roos et al. 2018), yet the relationship between prescribed fire in grasslands and wild bees is understudied. Fire reduces shrub encroachment, releases soil nutrients, and increases light availability, all of which boost forb abundance and diversity and increase blooms (Potts et al. 2003; Wagenius et al. 2020; Richardson and Wagenius 2022), which are expected to benefit pollinator communities (Nuland et al. 2013; Mola and Williams 2018; Gelles et al. 2023). However, fire may also have direct negative impacts on bees (i.e., mortality or reduced preference for recently burned habitats) or could disrupt floral resources during the critical spring queen nest searching phase (Mola and Williams 2018). Despite the widespread use of fire as a management tool, only a few studies have examined prescribed fire’s effects on prairie bee communities and the studies report conflicting results (Tai et al. 2022; Bruninga-Socolar et al. 2022; Leone et al. 2022).

To better understand how both tallgrass prairie restoration and management with prescribed fire affect wild bees, we measured the abundance and diversity of a focal genus, the bumble bees (*Bombus* spp.), at 32 sites in Southern Wisconsin over 2 years. We compared bumble bee abundance and diversity at restored grasslands that ranged in restoration intervention intensity: no intervention (unassisted recovery), low intervention (seed-only), and moderate intervention (seeded and periodically burned). Our study objectives were to determine whether (1) bumble bee abundance and diversity respond positively to assisted restoration of tallgrass prairie (e.g., low and moderate intervention restoration), and (2) bumble bee abundance and diversity differ between low-intensity (seeded only) and moderate-intensity (seeded followed by prescribed fire management) restorations.

## Methods

### Study Sites

We sampled bumble bees at 11 sites in 2018 and an additional 21 sites in 2019, totaling 32 study sites across Southern Wisconsin, USA. All study sites were former agricultural lands now managed under agreements with the U.S. Department of Agriculture (USDA) Natural Resources Conservation Service (NRCS) as part of the Wetland Reserve Easement (WRE) program (Fig. S1).

Using NRCS-WRE restoration and management records, we categorized sites into three intensity levels of restoration intervention (Fig. S2): (1) No intervention (*n* = 6): upland grassland left to naturally regenerate following cessation of agricultural activities; (2) Low intervention (*n* = 13): initially seeded with native prairie plants and not subsequently managed using fire; and (3) Moderate intervention (*n* = 13): initially seeded and periodically managed using prescribed fire after restoration. For additional details on restoration methods and the plant communities in these categories, see Appendix A and McFarlane et al. 2023.

Low intervention sites were typically seeded with native prairie plants 1 - 2 years after enrollment into the WRE program with no additional management thereafter. Using available management records, we categorized “moderate intervention” sites as those that were burned at least once following restoration seeding of native prairie. These sites varied in fire regime; specifically burn frequency (e.g., once in 10 years vs. every 3 years), time since last burn, and seasonality. No sites surveyed in this study were considered “high intervention intensity”, i.e., a combination of soil amendments, repeated seeding, regular prescribed fire, and mechanical or chemical removal of shrubs and invasive plants. Total site size ranged from 1.6 to 810 ha (median: 31.5 ha), and time since enrollment in the WRE program ranged from 3 - 22 years (median: 10 years). Sites ranged in amount of semi-natural habitat in the landscape (6.9% - 75.5% in 1.5-km buffer; IQR: 28.7 - 54.1, median: 40.9%), though they were not selected to do so during initial study design.

### Bumble Bee Community Sampling

We measured bumble bee abundance and species richness at each site from 16 July - 4 September in 2018 and 3 June - 27 August in 2019. Sites were visited twice in 2018 and three times in 2019, every 3 - 5 weeks. Surveys were conducted on sunny to partly cloudy days between 09:00 and 17:30, with temperatures between 16 - 35 °C and wind speeds below ~ 7 m/s. We used non-lethal hand-netting catch-and-release survey protocols based on the U.S. Fish and Wildlife Service’s Survey Protocols for the Rusty Patched Bumble Bee (US Fish and Wildlife Services 2017) under Endangered Species Recovery Permit Number: ES06130D. Two types of timed surveys were conducted at each visit to control for observation effort and bias: transect-based and meandering surveys.

### Transect Surveys

We sampled bumble bee abundance and species richness along three 90-m transects per site during each visit. Observers walked each transect at a slow pace, alternating 2-min walking with 1-min stationary observations, for a total of 10 person-minutes of active observation per transect, or 30 person-minutes across all transects at a site per visit. Bumble bees observed within a 2-m area (length of insect net and observer’s arm) in front of and on both sides of the observer were hand-netted, identified to species (Table S3), and released on the transect from which they were caught. If identification and handling exceeded 2-min (less time in high heat), bees were chilled on ice to take pictures for later identification. While handling bees, we paused the timer to ensure the duration of each survey represents only active observation time. Several bumble bee species in our region could not be reliably distinguished in the field using catch-and-release methods. As a result, we combined hard to identify species into species complexes: *B. vagans* and *B. sandersoni* (*B*. vagans-sandersoni) and *B. auricomus* and *B. pensylvanicus* (*B*. auricomus-pensylvanicus; Table S3).

To account for systematic biases related to observer, space, and time, we randomized the order of site visits, surveyed only during optimal weather conditions, and the same two observers caught and identified bumble bees in both years. From these standardized methods, we calculated bumble bee abundance as the number of bees caught per person-hour per visit (bees/hour/visit) for the transect surveys.

### Meandering Surveys

To better estimate bumble bee species richness and diversity (Roswell et al. 2021) we conducted one 25 person-minute meandering survey per visit, targeting areas near the transects with high floral density, when available, to capture patchiness in flower distribution that may have been missed with the transect approach. The same catch and release hand-netting methods described above were used to capture and identify bumble bees, and up to 3-min of observation time was allowed at a single high floral density patch.

We estimated bumble bee diversity using the inverse Simpson’s diversity index (1/D). To do so, we combined bumble bee species data from the transect and meandering surveys and summed the abundance of each species across all visits within a year to obtain a site-by-species matrix with cells representing total abundance of each bumble bee species per site. The inverse Simpson’s diversity index (1/D) was calculated using *diversity*(index = “invsimpson”) in the *vegan* package in R (v3.6.1; Oksanen et al. 2025), since interpretation of this index is more intuitive (e.g., higher values indicate greater diversity, probability two individuals in a sample are the same species; Roswell et al. 2021).

### Flowering Plant Abundance Estimation

Bumble bee responses to restoration intervention intensity are expected to respond to the abundance of floral resources since bumble bees are obligatory flower visitors. To characterize site-level flowering plant abundance, we recorded the presence of each flowering species in four 1 x 1-m quadrats spaced every 10-m along each of the three 90-m transects used for bumble bee surveys (96 total quadrats per site). For this analysis, we calculated flowering plant abundance as the number of quadrats in which each flowering plant species occurred, summing those counts across all species and quadrats for a given visit at each site. Thus, abundance represents the total number of quadrat-level species occurrences at a site and not simply the number of quadrats with flowers present. Multiple species in a single quadrat contribute multiple occurrences (e.g., two species in one quadrat adds 2 to abundance). Detailed methods and a broader analysis of the plant community can be found in (McFarlane et al. 2023).

### Statistical Analyses

#### Differences in bumble bee abundance and diversity across restoration intervention intensities

We used generalized linear mixed models (GLMMs) to test whether bumble bee abundance and diversity differed across levels of restoration intervention intensity. We hypothesized that bumble bee abundance and diversity would increase with restoration intervention intensity (i.e., no intervention < low < moderate). Our models included restoration intervention, floral abundance, and landscape as main fixed effects. To account for repeated measures within a year and interannual variation, site, visit, and year were included as a random effects (Table S1). Given that landscape context can influence bee responses to local management (e.g., Tscharntke and Greiler 2003; Albrecht et al. 2007; Grab et al. 2018) we initially explored the possibility of including the interactions between a landscape covariate (% semi-natural habitat (SNH); sum of grassland, forest, wetland) in a 1.5-km buffer around each site, and both restoration intervention and floral abundance. Due to high multicollinearity and convergence issues, we used a stepwise model selection approach to compare candidate GLMMs varying in main effects, interaction terms, and error structure (Table S1).

All GLMMs were fit using the *glmmTMB* package (v1.1.10; Brooks et al. 2017) in R (v4.4.1; R Core Team 2024). Model diagnostics, including for overdispersion, and residuals normality, and heteroscedasticity, were performed using the *DHARMa* package (v0.4.7; Hartig 2016). To identify the best fitting and most parsimonious model, we evaluated model fit using *check_collinearity*() and *r2*() from the *performance* package (v0.14.0; Lüdecke et al. 2021), and compared models using *AIC*() from the *stats* package in R. We removed the interaction terms involving SNH from the final model given high multicollinearity (interactions: VIF > 10) and their inclusion increased model complexity without improving explanatory power (Table S1; Fig. S3).

The final model used for inference included fixed effects of restoration intervention, floral abundance, and SNH. For this model, we conducted analysis of variance using *Anova*() in the *car* package (v3.1.3; Fox et al. 2024), applying Type II sums of squares to account for uneven sample sizes across restoration categories and years. When overall significant effects were detected (alpha = 0.05), we used the *emmeans* package (v1.10.5; Lenth 2025) to calculate estimated marginal means among restoration categories. Standardized effect sizes were calculated using *model_parameters*(standardize = “basic”) from the *parameters* package (v0.27.0; Lüdecke et al. 2019).

#### Differences in bumble bee community composition across restoration intervention intensities

To evaluate differences in bumble bee community composition among restoration categories, we focused our analysis on the core species community. That is, species consistently present across most sites and likely driving community-level responses. Species contributing <5% of the total abundance were excluded to minimize the disproportionate effect of rare species on pairwise dissimilarity among sites (McCune and Grace 2002; Clarke et al. 2006). This removed *B. affinis, B. borealis, B. citrinus*, and *B. fervidus*. Bray-Curtis dissimilarities were calculated from a pairwise distance matrix calculated using total site-level abundance of each bumble bee species, summed across all visits over a season. Prior to ordination, data were square root transformed to reduce the influence of dominant species (Legendre and Gallagher 2001). One site was excluded from this analysis since no bumble bees were observed.

We used permutational multivariate analysis of variance (PERMANOVA; Anderson 2006) to test for differences in bumble bee community composition across restoration category and year. The *adonis2*() function in the *vegan* package was used to sequentially assess marginal effects of each term, with residuals permuted under a reduced model (Freedman and Lane 1983). PERMANOVA significance was assessed with 9,999 permutations. Pairwise comparison among restoration intervention intensity were conducted using *pairwise*.*adonis*(sim.function = “vegdist”, sim.method = “bray”) from the *pairwiseAdonis* package (v0.4.1; Martinez Arbizu 2020).

To assess whether differences in composition were due to variation in community dispersion rather than centroid position, we tested for homogeneity of multivariate dispersions across categories using *betadisper*() followed by *permutest*() in *vegan* (Anderson 2006).We visualized community patterns from PERMANOVA using nonmetric multidimensional scaling (NMDS) with *metaMDS*() also in *vegan* and exported scores to be plotted with *ggplot2*. We report permutation-based *P*-values since they provide an exact test of our null hypotheses (Anderson et al. 2010).

## Results

We observed 1,727 individual bumble bees representing 10 species (Table S3), ranging from 1 to 173 individuals (1 to 8 species) per site in 2018, and 1 to 75 individuals (1 to 9 species) per site in 2019. The most common species cumulatively representing ~ 90% of all observed individuals included *B. impatiens, B. griseocollis, B. vagans-sandersoni, B. auricomus-pensylvanicus*, and *B. rufocinctus*. Uncommon species which represented less than 10% of all observed individuals included *B. affinis* (federally endangered), *B. bimaculatus, B. borealis, B. citrinus* (social parasite), and *B. fervidus* (state species of concern).

### Bumble bee abundance and diversity at sites with varying restoration intervention intensity

Bumble bee abundance did not significantly differ with restoration intervention intensity (GLMM: χ^2^ = 3.87, *P* = 0.145; Fig. 1A). However, we still evaluated pairwise differences between categories of restoration intervention intensity using estimated marginal mean contrasts and Tukey’s HSD. At sites that underwent any restoration intervention (e.g., low and moderate), bumble bee abundance was 2.7x higher than sites with no intervention (*P* = 0.060). Bumble bees were 3x and 2.4x higher at moderate and low intervention intensity sites, respectively, when compared to sites with no intervention, though these increases were not significant after adjusting for multiple comparisons with Tukey’s HSD (moderate: *P* = 0.121, low: *P* = 0.238). Bumble bee abundance was only 22% higher when comparing moderate to low intervention intensity sites (*P* = 0.819). Bumble bee abundance significantly increased with higher floral abundance (GLMM: χ^2^ = 26.36, *P* < 0.0001; Fig. S3A) but not with the amount of semi-natural habitat in the landscape (GLMM: χ^2^ = 0.12, *P* = 0.726; Fig. S3B).

**Figure 1.**
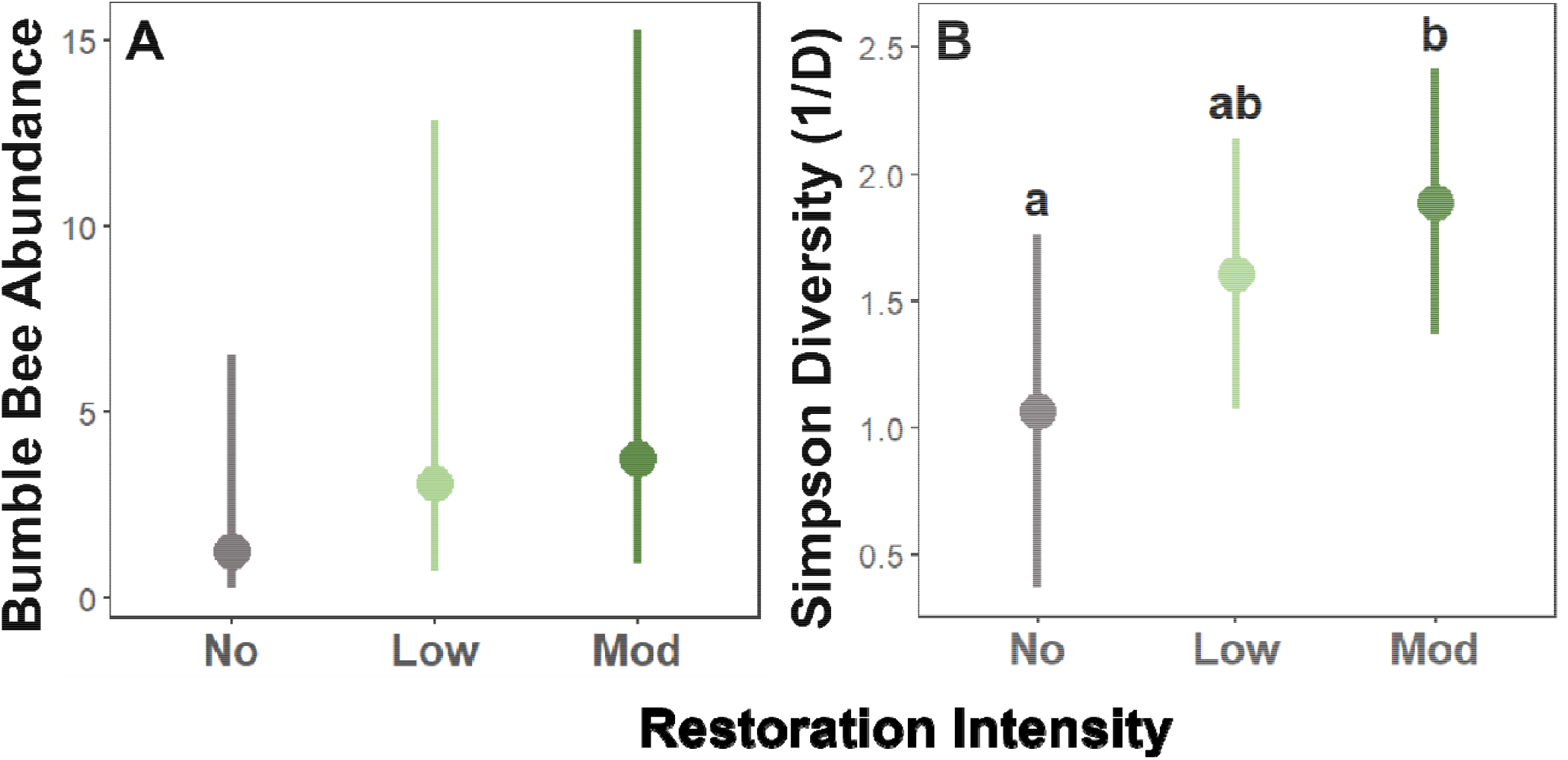
This figure shows the estimated marginal means (± 95% CI) of (A) bumble bee abundance and (B) inverse Simpson’s diversity (1/D) across levels of prairie restoration intervention intensity (partial residual effects after accounting for other fixed and random effects in the model; Table 1). Compared to no restoration intervention, bumble bee abundance had an increasing trend with restoration intervention, though it was not significant (Type II Wald χ^2^ = 3.87, df = 2, P = 0.145), and inverse Simpson’s diversity did significantly increase (Type II Wald χ^2^ = 6.42, df = 2, P = 0.040). Different letters indicate statistically significant differences between restoration intervention intensities based on pairwise comparisons using Tukey-adjusted p-values (p-value < 0.05; *emmeans*).

Bumble bee diversity significantly increased with restoration intervention intensity (GLMM: χ^2^ = 6.42, *P* = 0.040; Fig. 1B). Bumble bee diversity was 2x higher at sites that underwent any restoration intervention compared to those that had no restoration intervention. Diversity increased the most at moderate intervention sites; it was 2.3x higher than no intervention. Low intervention sites had 72.2% higher diversity than no intervention and there was only a 33% increase between low and moderate intervention sites. Diversity also significantly increased with floral abundance (GLMM: χ^2^ = 7.85, *P* < 0.001; Fig. S3C) but not the amount of semi-natural habitat in the landscape (GLMM: χ^2^ = 0.0736, *P* = 0.786; Fig. S3D).

### Bumble bee community composition across sites with varying restoration intervention intensity

In this study, bumble bee community composition was defined as the identity and relative abundance of the *Bombus* species that accounted for >95% of total observations. We found that overall (core) community composition differed significantly across types of restoration intervention intensities (PERMANOVA: Pseudo-F_2,37_ = 3.31, *P* = 0.0005, stress = 0.19; Fig. 2). Pairwise PERMANOVA demonstrated that sites with no restoration intervention were most different from moderate intervention (Pseudo-F = 5.35, *P* = 0.003). But low intervention sites differed only slightly from sites with no intervention (Pseudo-F = 2.87, *P* = 0.078) and did not differ from moderate intervention sites (Pseudo-F = 1.67, *P* = 0.375). The difference in community composition between no and moderate intervention sites is likely driven by the two dominant species, *B. impatiens* and *B. griseocollis*, becoming less dominant. *Bombus impatiens* and *B. griseocollis* made up 82% of bees observed at no intervention sites and decreased in relative abundance to comprise 76% of observations at moderate intervention sites (Table S2, Fig. S4). The less frequently, but still relatively commonly encountered species *B. auricomus/pensylvanicus, B. vagans/sandersoni, B. rufocinctus*, and B. *bimaculatus* all increased with restoration intervention intensity and, in some cases, increased nearly an order of magnitude (Table S2). Together, this evidence suggests that the significant differences reflect shifts in abundance of the core bumble bee community across types of restoration intervention (Fig. S4). Though not included in the formal community analysis because of their rarity, it is notable that species of conservation concern (*B. affinis, B. fervidus*) were only observed at sites with active restoration interventions (Fig. S4).

**Figure 2.**
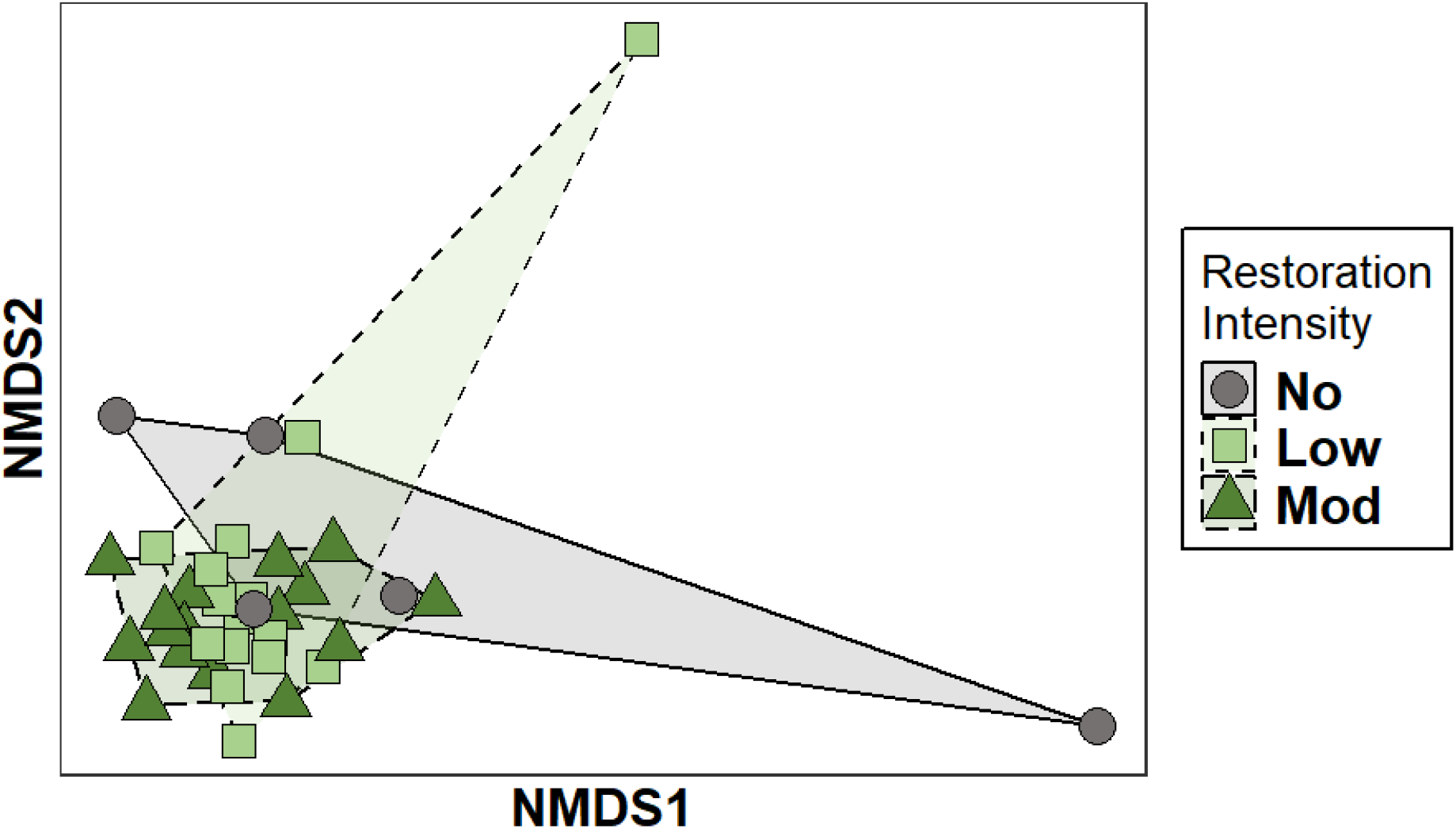
Nonmetric multidimensional scaling (NMDS) ordination that represents bumble bee community composition in two dimensions. The NMDS is based on Bray-Curtis dissimilarities of low and moderate (“mod”) restoration intervention intensity compared to no restoration intervention (PERMANOVA, *Pseudo-F*_2,36_ = 2.36, *P* = 0.009; Stress = 0.128; Table 2). Each point represents a site and shaded regions are convex hulls around points of each restoration intensity category. Shapes and colors correspond to restoration categories.

### Flowering plant abundance across sites with varying restoration intervention intensity

We observed 7,612 flowering plant occurrences representing 133 taxa (Table S4). Abundance ranged from 8 to 398 occurrences (4 to 27 species) per site in 2018, and 1 to 509 occurrences (1 to 32 species) per site in 2019. Twenty-four species made up 80% of all flowering plant occurrences (Table S4). Of these species, 13 were observed being used by *Bombus affinis* in this study and elsewhere in Wisconsin (Wolf et al. 2022).

Floral abundance was significantly influenced by restoration intervention intensity (GLMM: χ^2^ = 11.38, *P* = 0.003; Fig. 3, Table S1). Moreover, there was a significant interaction with the amount of semi-natural habitat in the surrounding landscape (GLMM: χ^2^ = 6.94, *P* = 0.031; Fig. S5), though landscape did not have its own effect on floral abundance (GLMM: χ^2^ = 0.35, *P* = 0.552). Floral abundance increased 73.5% at low intervention sites compared to no intervention (when estimating marginal means at the median value of semi-natural amount ~42%; *P* = 0.200). There were 3x more flowers at moderate intervention sites relative to no intervention sites (*P* = 0.001) and ~74% more compared to low intervention (*P* = 0.007). The interaction effect between restoration intervention intensity and semi-natural habitat in the landscape was apparent at no intervention sites, where floral abundance increased 4.6x over the range of semi-natural habitat (*P* = 0.014). Floral abundance at low intervention sites decreased by ~36% over this same range and moderate intervention sites increased by just 15%, though neither of these trends were significant (*P* = 0.281 and *P* = 0.725, respectively).

**Figure 3.**
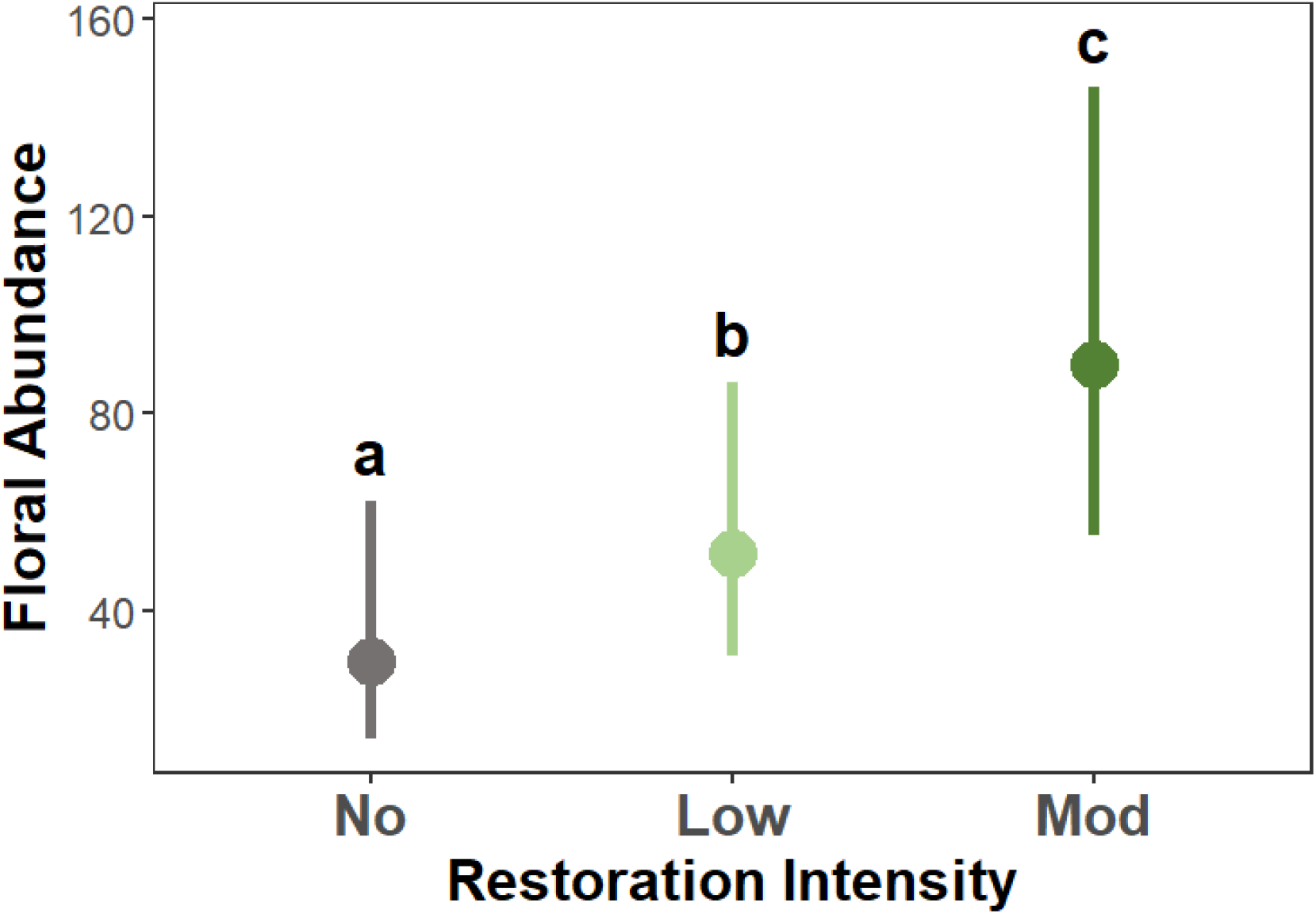
Estimated marginal mean response (± 95% CI) of floral abundance across prairie restoration intervention intensities (partial residual effects after accounting for other fixed and random effects in the model; Table S1). Floral abundance, or the number of flower occurrences, significantly increased with moderate restoration intervention intensity when compared to no and low restoration intervention (Type II Wald χ^2^ = 11.52, df = 2, P = 0.003). Different letters indicate statistically significant differences between restoration intensities based on pairwise comparisons using Tukey-adjusted p-values (p-value < 0.05; *emmeans*).

## Discussion

Restoration and management of native habitats may be vital to counteracting bee population declines. This study evaluated how bumble bee populations respond to varying intensities of grassland restoration interventions. We found that actively restored tallgrass prairies, which were either seeded with native plants or both seeded and managed with fire, had nearly three times as many bumble bees and twice the diversity than sites that were left to passively regenerate following cessation of agricultural activities (i.e., old fields or no restoration intervention). These increases in bumble bee abundance were associated with higher floral abundance also present at more actively managed sites. Additionally, sites with the highest intervention intensity (seeded and managed with fire) had different composition of bumble bee communities relative to no intervention sites. Our findings highlight that site-level habitat restoration can be important for bumble bee conservation and reinforce other studies showing positive effects of grassland restoration for wild bees (Tonietto et al. 2017; Denning and Foster 2018; Lane et al. 2020). Although many wild bee species have declined in population size and geographic range due to habitat loss, the positive effects of tallgrass prairie restoration on bumble bees provide land managers opportunities to use plant community restoration to support pollinators like bumble bees.

Implementing restoration interventions that increase overall bumble bee population size can have benefits across species. Of the approximately 20 bumble bee species that occur in Wisconsin, we observed 10 in this study. However, the community was dominated by just two species, *B. impatiens* and *B. griseocollis*, which together accounted for over half of all observations. Despite this dominance, abundance across nearly all species doubled with each increase in restoration intervention intensity (though this was not statistically significant). Increases in diversity followed a similar pattern. Diversity nearly doubled between no and moderate intervention sites. Since Inverse Simpson’s Diversity index gives greater weight to common species, this change primarily reflects shifts among dominant and moderately common species rather than the addition of rare species. Indeed, rare species comprised less than 5% of all observations and were not included in statistical analyses. These patterns are consistent with expectations from species accumulation curves (Roswell et al. 2021), which indicate that as abundance increases, so does the number of observed species. Thus, our results suggest that differences in diversity largely reflect changes in abundance and relative dominance of a core community of species across restoration types. Although rare species were not included in formal analyses, their occurrence patterns are noteworthy. Species of conservation concern (*B. affinis, B. fervidus*) were only observed at sites with active restoration interventions. Moreover, observations of *B. affinis* and *B. fervidus* doubled between low to moderate intervention intensity, which suggests that management may benefit vulnerable or declining bumble bee species.

The positive effects of tallgrass prairie restoration on bumble bees may be explained by several mechanisms. First, restoration via seeding of native prairie plants increased the amount and quality of floral resources, both of which are important components of nutritious food for bees (Vaudo et al. 2015). We observed increases in floral abundance, and concurrent work in this system showed increases in floristic quality and the number of native plant species (i.e., species richness), though the latter effect was not statistically significant (McFarlane et al. 2023). Our findings align with previous studies demonstrating that greater flower abundance and species richness support higher pollinator abundance and species richness (e.g., Blaauw and Isaacs 2014; Lane et al. 2020). Second, restored sites contained flower species preferred by bumble bees of conservation concern, while no intervention sites were dominated by non-native plants that are used but not necessarily preferred (Williams et al. 2011; Seitz et al. 2020). Therefore, increases in bumble bee responses may primarily reflect greater overall floral abundance, with additional benefits arising from increases in preferred floral resources, including those used by the endangered *Bombus affinis*, which we observed more frequently at sites containing its known forage plants (Harmon-Threatt and Hendrix 2015; Novotny et al. 2021; Wolf et al. 2022). However, higher bumble bee abundance at actively restored sites may result from either concentration effects (where bees move into restorations from the surrounding landscape to forage) or actual population growth supported by increased floral resources (Tonietto and Larkin 2018; Lane et al. 2022). Although our study was not designed to distinguish these mechanisms, the results nonetheless suggest that prairie restorations support higher bumble bee abundance and diversity.

Among sites where restoration intervention occurred, we expected large differences between non-burned (low intensity) and burned (moderate intensity) sites due to fire-induced increases in floral abundance and diversity (Nuland et al. 2013; Mola and Williams 2018). However, we did not observe large differences between low and moderate intensity sites. One possibility for this is that fire regimes (i.e., time since last burn, frequency of fire management) were highly variable among moderate intervention sites. Based on available management records, we categorized “burned” sites as those that were burned at least once post-restoration seeding. Therefore, burn frequency varied among sites in the “burned” category (e.g., once in 10 years vs. every 3 years), time since last burn, and seasonality, factors that can substantially influence plant communities (Knapp et al. 2009; Veldman et al. 2014; Decker and Harmon-Threatt 2019). In particular, floral abundance is known to be generally higher shortly after prescribed fires and then declines as grasses become more dominant (Vogel et al. 2010; Decker and Harmon-Threatt 2019). Despite these limitations, these prairie restorations represent typical variability in fire management practices used by land stewards in this region. Our findings support recent studies showing that prescribed fire was not associated with lower bumble bee abundance or diversity in tallgrass prairies (Tai et al. 2022; Bruninga-Socolar et al. 2022), a pattern also observed for other pollinator taxa and ecosystems (Glenny et al. 2022).

In addition to the local-scale environment, we examined whether patterns of bumble bees communities and floral resources were affected by characteristics of the surrounding landscape (e.g., Hines and Hendrix 2005; Mallinger et al. 2016; Spiesman et al. 2019; Griffin et al. 2021). We expected that greater amounts of semi-natural habitat in the landscape would increase bumble bee abundance and diversity at both old fields and actively restored prairies, and that floral abundance would follow similar patterns. However, we found no evidence of landscape influence on bumble bee communities nor an interaction with restoration intervention intensity, consistent with other recent studies (e.g., Lane et al. 2020; Leone et al. 2022). We found that for old fields only, floral abundance was greater in landscapes with more natural habitat. This suggests that in the absence of active seeding, landscape level seed sources are important for floral resource establishment. Though we did not choose study sites to specifically explore gradients in landscape characteristics, our sites were located across a broad range of semi-natural habitat in the landscape, from highly agricultural (6.9 % of landscape in semi-natural habitat) to mostly natural habitat (75.5 % SNH; median: 40.9 %), and this gradient was represented within all categories of restoration intervention intensity. While our study does not support the idea that broad-scale landscape characteristics influence bumble bee communities in prairie restorations, a randomized selection of restorations across a landscape gradient would be needed to more fully evaluate whether bees are affected by the interaction between local-scale prairie restoration and the amount of semi-natural habitat in the landscape. Instead, we found strong evidence that local-scale restoration of plant communities via native tallgrass prairie seeding and fire may be, in this landscape context, relatively more important for bumble bee conservation.

While we focus on restoration interventions of seeding and prescribed fire, restoration efforts may have influenced bumble bee communities in other ways, and through changes in unmeasured environmental attributes. For example, our surveys focused on bumble bee workers foraging during the summer, but restoration may also affect other stages of the bumble bee life cycle (e.g., queens/gynes establishing nests or overwintering) via alterations to habitat structure (e.g., amount of bare ground, Spiesman et al. 2019). Because these features are likely to affect nest establishment and overwinter survival (reviewed in Harmon-Threatt 2020), research could incorporate measurements of habitat structure or grass tussocks (Buckles and Harmon-Threatt 2019). Additionally, distinguishing among bumble bee species with differing nesting ecologies (e.g., above versus below ground; Decker and Harmon-Threatt 2019) could provide a more comprehensive understanding of how prairie restoration and management affect these pollinators.

Our findings contribute to growing evidence that active grassland restoration is an effective conservation practice for bumble bees, particularly in terms of increasing local abundance and diversity. It is of conservation interest that the endangered *B. affinis* was observed at grassland sites with active restoration interventions, regardless of management history. Although fire regime (e.g., intensity, frequency, etc., rather than burned/not-burned) should be explored to better understand the effects of prescribed fire on bumble bees, our study suggests that management using prescribed fire does not negatively impact bumble bees. In fact, bumble bee abundance and diversity increase with intervention intensity that included fire. Less common species, including species of concern like *B. affinis*, became more frequent with increasing intervention intensity. Overall, these positive effects of tallgrass prairie restoration on bumble bee communities can give land managers confidence that active management across a range of landscape contexts is an effective practice to support insect pollinator conservation. These results reinforce that biodiversity and population decline from habitat loss and fragmentation may be mitigated by habitat restoration.

## Supporting information

Associated Data

Rendered R Quarto Document

## Acknowledgements

This research would not have been possible without USDA-NRCS state office staff: Greg Kidd, Robert Geitner, Stephaney Olson, Gretchen Skudlarczyk, Tally Hamilton, and staff at NRCS field offices across southern Wisconsin. Thank you to the current land stewards: Dane County Parks Department (Lars Higdon), Wisconsin Department of Natural Resources (Julie Widholm, Nathan Holoubek, Jason Cotter), countless private landowners enrolled in the USDA-NRCS Wetland Reserve Easement program, and the generational land stewards of the Ho-Chunk, whose ancestral lands are the modern study area. Eternal gratitude to our undergraduate field research assistants: Audrey Spiegelhoff and Grant Witynski. Microsoft Word and Google Docs were used to improve spelling, grammar, and general editing. Both ChatGPT and Claude.AI were used to troubleshoot and edit R code for reproducibility. Data files are available via the Dryad Digital Repository https://doi.org/10.5061/dryad.rxwdbrvq5 (Kochanski et al. 2025) and can be accessed for review using the following link: http://datadryad.org/share/LINK_NOT_FOR_PUBLICATION/dFm5P6n87ix94PZBAJvRnvYudQBCET4MPx9GLYX8Pys

---

**Table 1.**
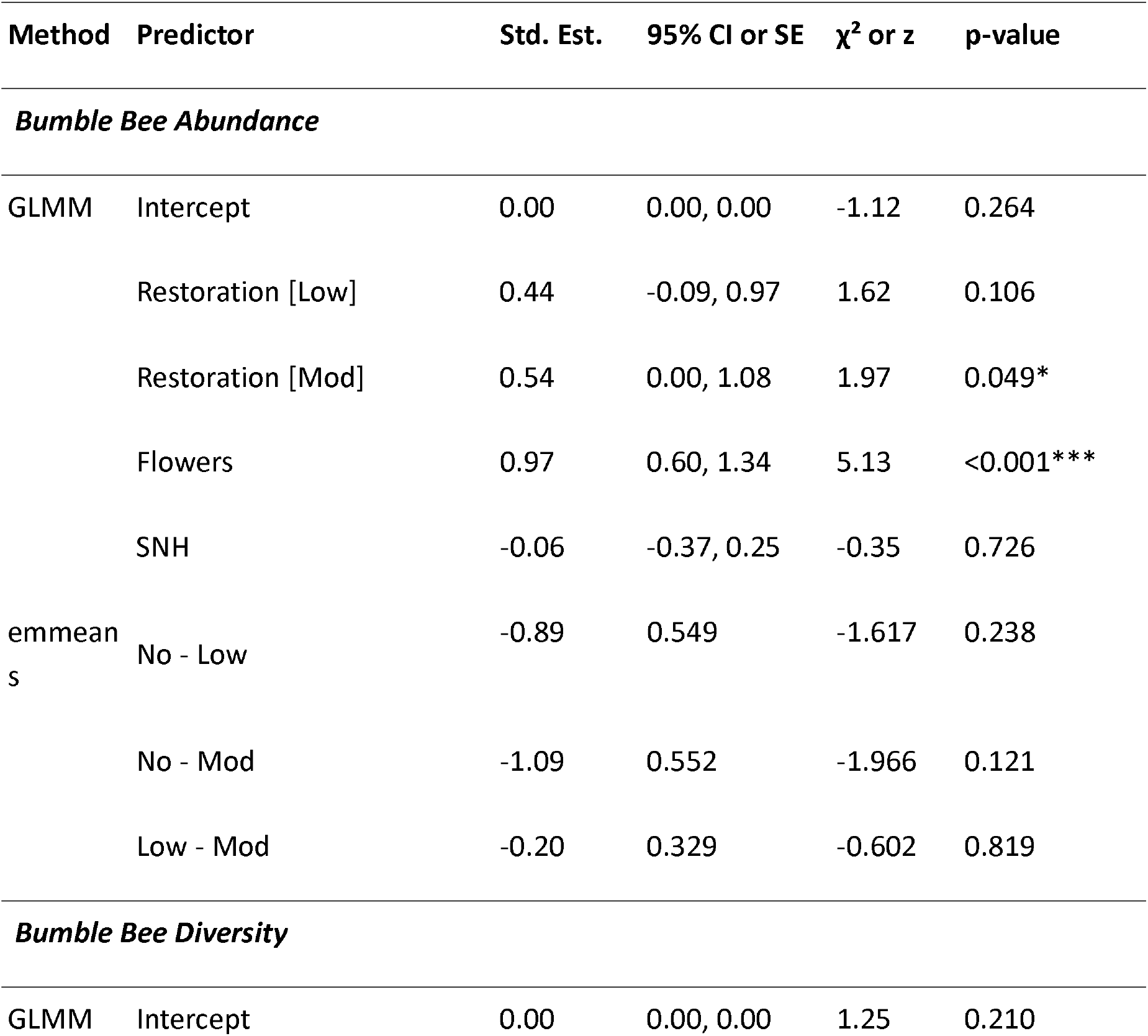
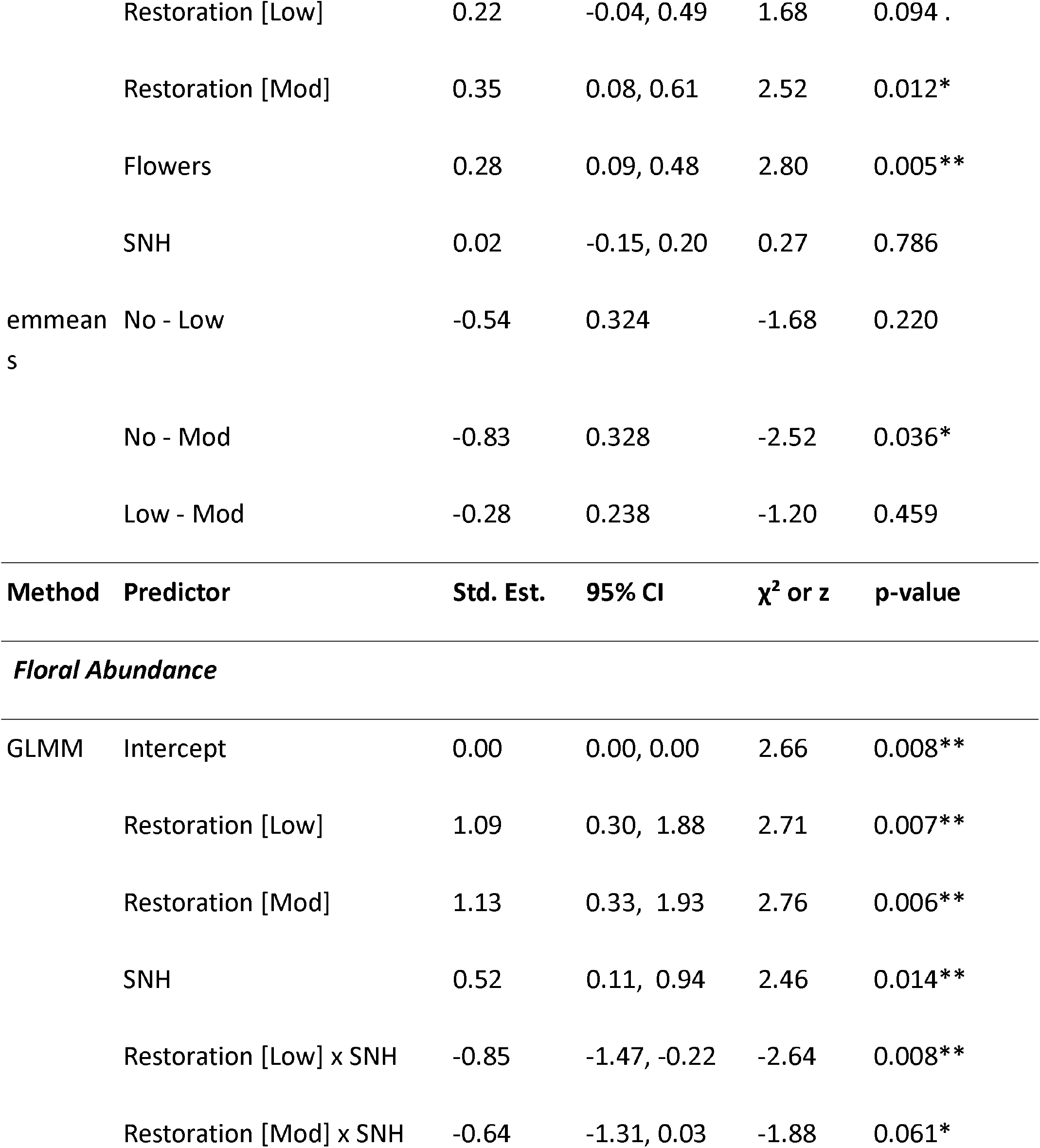
Summary of model results and post-hoc comparisons for bumble bee abundance, inverse Simpson’s diversity, and floral abundance. Generalized linear mixed models (GLMMs) with Type II Wald χ^2^ tests were used for overall fixed effects, with estimated marginal means and pairwise comparison using Tukey-adjusted *z*-tests. Standardized estimates and 95% CI were generated from *parameters* package in R (v0.27.0; Lüdecke et al., 2019). Significant predictors are indicated by asterisks: p < 0.05 (*), p < 0.01 (**), and p < 0.001 (***).

**Table 2.** Summary of permutation-based multivariate analysis of variance (PERMANOVA) results testing the effects of restoration on bumble bee community composition. Overall results were generated using *vegan::adonis2*(permutations = 999, method = “bray”, by = “margin”, model = “reduced”). Pairwise comparisons between restoration intervention categories are from *pairwise*.*adonis*(), which adjusts p-values for multiple comparisons using the Bonferroni method. Significant predictors are indicated by asterisks: p < 0.05 (*), p < 0.01 (**), and p < 0.001 (***).

| Predictor | df | Sums of Sqs | $R^2$ | pseudo-F | *p* | *p-adjusted* |
| --- | --- | --- | --- | --- | --- | --- |
| Restoration | 2 | 0.58 | 0.12 | 2.36 | 0.009** | - |
| Flowers | 1 | 0.09 | 0.02 | 0.77 | 0.574 | - |
| SNH | 1 | 0.05 | 0.01 | 0.39 | 0.858 | - |
| Year | 1 | 0.08 | 0.02 | 0.63 | 0.694 | - |
| No - Low | 1 | 0.39 | 0.12 | 2.59 | 0.030* | 0.090 |
| No - Mod | 1 | 0.58 | 0.19 | 4.70 | 0.004** | 0.012* |
| Low - Mod | 1 | 0.16 | 0.05 | 1.67 | 0.131 | 0.393 |

## Notes

https://datadryad.org/share/LINK_NOT_FOR_PUBLICATION/dFm5P6n87ix94PZBAJvRnvYudQBCET4MPx9GLYX8Pys

### Competing Interest Statement

The authors have declared no competing interest.

