## Supplementary material for "Bumble Bee Abundance and Diversity Increase with Intensity of Tallgrass Prairie Restoration Intervention": Rendered R Quarto Document

Restoration intervention intensity

Jade M. Kochanski

This documents contains final analyses for the publication/manuscript of Jade Kochanski’s 2018 - 2019 Master’s research examining increases in bumble bee abundance and diversity with tallgrass prairie restoration in Wisconsin, USA. Analyses and code finalized October 2025.

### Load packages

```
# general
library(tidyverse) # general commands, data wrangling
library(here) # call data regardless of file paths


# analyses
library(pairwiseAdonis) # pairwise PERMANOVA
library(glmmTMB) # generalized mixed models
library(DHARMa) # check model assumptions
library(ggeffects) # predicting model values
library(emmeans) # also for model predictions
library(car) # statistical analysis
library(performance) # multicollinearity
library(parameters) # effect sizes
library(partR2) # partial R2

# graphing packages
library(ggpubr) # graphing
library(grid) # graphing
library(gridExtra) # graphing
library(cowplot) # graphing
```

### Load data

note: for using the library(here), change the path to the folder name in your working directory in which the csv data files are located.

```
# visit-level bumble bee responses and associated predictors
bombus_responses <- readr::read_csv(
  here("data_output", "bombus_responses.csv")
)  %>%
  # specify level order for correct reference level
  mutate(
    RestorationCategory = factor(
      RestorationCategory,
      levels = c("No", "Low", "Mod"))
    )
Rows: 96 Columns: 10
── Column specification ────────────────────────────────────────────────────────
Delimiter: ","
chr (4): EasementID, Visit, RestorationCategory, County
dbl (6): Year, TotalBees, invSimp, SemiNatural, TotalFlowers, CenteredFlowers

ℹ Use `spec()` to retrieve the full column specification for this data.
ℹ Specify the column types or set `show_col_types = FALSE` to quiet this message.
# visit-level floral observation data
floral_incidence <- readr::read_csv(
  here("data_output", "floral_incidence.csv")
) %>%
  # specify level order for correct reference level
  mutate(
    RestorationCategory = factor(
      RestorationCategory,
      levels = c("No", "Low", "Mod"))
    )
Rows: 97 Columns: 6
── Column specification ────────────────────────────────────────────────────────
Delimiter: ","
chr (3): EasementID, Visit, RestorationCategory
dbl (3): Year, TotalFlowers, SemiNatural

ℹ Use `spec()` to retrieve the full column specification for this data.
ℹ Specify the column types or set `show_col_types = FALSE` to quiet this message.
# site by species matrix for community composition analysis
community_matrix <- readr::read_csv(
  here("data_output", "community_matrix.csv")
)
Rows: 37 Columns: 6
── Column specification ────────────────────────────────────────────────────────
Delimiter: ","
dbl (6): BOMAUR, BOMBIM, BOMGRI, BOMIMP, BOMRUF, VAGSAN

ℹ Use `spec()` to retrieve the full column specification for this data.
ℹ Specify the column types or set `show_col_types = FALSE` to quiet this message.
# environmental variables corresponding with site x spp matrix observations
community_env <- readr::read_csv(
  here("data_output", "community_env.csv")
) %>%
  # specify level order for correct reference level
  mutate(
    RestorationCategory = factor(
      RestorationCategory,
      levels = c("No", "Low", "Mod"))
    )
Rows: 37 Columns: 5
── Column specification ────────────────────────────────────────────────────────
Delimiter: ","
chr (2): EasementID, RestorationCategory
dbl (3): Year, SemiNatural, TotalFlowers

ℹ Use `spec()` to retrieve the full column specification for this data.
ℹ Specify the column types or set `show_col_types = FALSE` to quiet this message.
```

### Statistical analyses

##### Difference in floral abundance between sites w/ different restoration and management history

```
m1fa <- glmmTMB(TotalFlowers ~ RestorationCategory*SemiNatural +
                      (1 | Visit) +
                      (1| EasementID),
                    family = nbinom1,
                    data= floral_incidence)

summary(m1fa)
 Family: nbinom1  ( log )
Formula:
TotalFlowers ~ RestorationCategory * SemiNatural + (1 | Visit) +
    (1 | EasementID)
Data: floral_incidence

      AIC       BIC    logLik -2*log(L)  df.resid
   1029.7    1052.8    -505.8    1011.7        88

Random effects:

Conditional model:
 Groups     Name        Variance Std.Dev.
 Visit      (Intercept) 0.1321   0.3635
 EasementID (Intercept) 0.0608   0.2466
Number of obs: 97, groups:  Visit, 3; EasementID, 32

Dispersion parameter for nbinom1 family (): 41.8

Conditional model:
                                   Estimate Std. Error z value Pr(>|z|)
(Intercept)                         2.09365    0.78679   2.661  0.00779 **
RestorationCategoryLow              2.20200    0.81331   2.707  0.00678 **
RestorationCategoryMod              2.26979    0.82371   2.756  0.00586 **
SemiNatural                         0.03049    0.01242   2.455  0.01408 *
RestorationCategoryLow:SemiNatural -0.03924    0.01489  -2.635  0.00841 **
RestorationCategoryMod:SemiNatural -0.02771    0.01477  -1.876  0.06069 .
---
Signif. codes:  0 '***' 0.001 '**' 0.01 '*' 0.05 '.' 0.1 ' ' 1
car::Anova(m1fa)
Analysis of Deviance Table (Type II Wald chisquare tests)

Response: TotalFlowers
                                  Chisq Df Pr(>Chisq)
RestorationCategory             11.3756  2   0.003387 **
SemiNatural                      0.3532  1   0.552315
RestorationCategory:SemiNatural  6.9436  2   0.031062 *
---
Signif. codes:  0 '***' 0.001 '**' 0.01 '*' 0.05 '.' 0.1 ' ' 1
# standardized estimates and transform to response scale
model_parameters(m1fa,
                 standardize = "basic")
Warning: Standardizing coefficients only works for fixed effects of the mixed
  model.
# Fixed Effects

Parameter                               | Std_Log-Mean |   SE |         95% CI
------------------------------------------------------------------------------
(Intercept)                             |         0.00 | 0.00 | [ 0.00,  0.00]
RestorationCategory [Low]               |         1.09 | 0.40 | [ 0.30,  1.88]
RestorationCategory [Mod]               |         1.13 | 0.41 | [ 0.33,  1.93]
SemiNatural                             |         0.52 | 0.21 | [ 0.11,  0.94]
RestorationCategory [Low] × SemiNatural |        -0.85 | 0.32 | [-1.47, -0.22]
RestorationCategory [Mod] × SemiNatural |        -0.64 | 0.34 | [-1.31,  0.03]

Parameter                               |     z |     p
-------------------------------------------------------
(Intercept)                             |  2.66 | 0.008
RestorationCategory [Low]               |  2.71 | 0.007
RestorationCategory [Mod]               |  2.76 | 0.006
SemiNatural                             |  2.46 | 0.014
RestorationCategory [Low] × SemiNatural | -2.64 | 0.008
RestorationCategory [Mod] × SemiNatural | -1.88 | 0.061

Uncertainty intervals (equal-tailed) and p-values (two-tailed) computed
  using a Wald z-distribution approximation.

The model has a log- or logit-link. Consider using `exponentiate =
  TRUE` to interpret coefficients as ratios.
```

##### Difference in bumble bee abundance and diversity between sites w/ different restoration and management history

###### Abundance

```
m3a <- glmmTMB(TotalBees ~ RestorationCategory +
                      TotalFlowers +
                      SemiNatural +
                      (1 | Year) +
                      (1 | Visit) +
                      (1| EasementID),
                    family = nbinom2,
                    data=bombus_responses)

summary(m3a)
 Family: nbinom2  ( log )
Formula:
TotalBees ~ RestorationCategory + TotalFlowers + SemiNatural +
    (1 | Year) + (1 | Visit) + (1 | EasementID)
Data: bombus_responses

      AIC       BIC    logLik -2*log(L)  df.resid
    425.2     448.3    -203.6     407.2        87

Random effects:

Conditional model:
 Groups     Name        Variance Std.Dev.
 Year       (Intercept) 0.3505   0.5920
 Visit      (Intercept) 0.7860   0.8866
 EasementID (Intercept) 0.1349   0.3673
Number of obs: 96, groups:  Year, 2; Visit, 3; EasementID, 32

Dispersion parameter for nbinom2 family (): 0.993

Conditional model:
                        Estimate Std. Error z value Pr(>|z|)
(Intercept)            -1.053258   0.943018  -1.117   0.2640
RestorationCategoryLow  0.887886   0.549136   1.617   0.1059
RestorationCategoryMod  1.085894   0.552260   1.966   0.0493 *
TotalFlowers            0.013955   0.002718   5.134 2.83e-07 ***
SemiNatural            -0.003231   0.009208  -0.351   0.7257
---
Signif. codes:  0 '***' 0.001 '**' 0.01 '*' 0.05 '.' 0.1 ' ' 1
car::Anova(m3a)
Analysis of Deviance Table (Type II Wald chisquare tests)

Response: TotalBees
                      Chisq Df Pr(>Chisq)
RestorationCategory  3.8662  2     0.1447
TotalFlowers        26.3610  1  2.832e-07 ***
SemiNatural          0.1231  1     0.7257
---
Signif. codes:  0 '***' 0.001 '**' 0.01 '*' 0.05 '.' 0.1 ' ' 1
# pairwise comparisons
# uses TukeyHSD for multiple comparisons
emmeans(m3a, pairwise ~ RestorationCategory)
$emmeans
 RestorationCategory  emmean    SE  df asymp.LCL asymp.UCL
 No                  -0.0833 0.837 Inf    -1.724      1.56
 Low                  0.8046 0.729 Inf    -0.624      2.23
 Mod                  1.0026 0.716 Inf    -0.401      2.41

Results are given on the log (not the response) scale.
Confidence level used: 0.95

$contrasts
 contrast  estimate    SE  df z.ratio p.value
 No - Low    -0.888 0.549 Inf  -1.617  0.2384
 No - Mod    -1.086 0.552 Inf  -1.966  0.1206
 Low - Mod   -0.198 0.329 Inf  -0.602  0.8192

Results are given on the log (not the response) scale.
P value adjustment: tukey method for comparing a family of 3 estimates
# standardized estimates and transform to response scale
model_parameters(m3a,
                 standardize = "basic")
Warning: Standardizing coefficients only works for fixed effects of the mixed
  model.
# Fixed Effects

Parameter                 | Std_Log-Mean |   SE |        95% CI |     z |      p
--------------------------------------------------------------------------------
(Intercept)               |         0.00 | 0.00 | [ 0.00, 0.00] | -1.12 | 0.264
RestorationCategory [Low] |         0.44 | 0.27 | [-0.09, 0.97] |  1.62 | 0.106
RestorationCategory [Mod] |         0.54 | 0.28 | [ 0.00, 1.08] |  1.97 | 0.049
TotalFlowers              |         0.97 | 0.19 | [ 0.60, 1.34] |  5.13 | < .001
SemiNatural               |        -0.06 | 0.16 | [-0.37, 0.25] | -0.35 | 0.726

Uncertainty intervals (equal-tailed) and p-values (two-tailed) computed
  using a Wald z-distribution approximation.
```

###### Diversity

```
m2d <- glmmTMB(invSimp ~ RestorationCategory +
                      TotalFlowers +
                      SemiNatural +
                      (1 | Visit) +
                      (1| EasementID),
                    family = gaussian,
                    data=bombus_responses)

summary(m2d)
 Family: gaussian  ( identity )
Formula:          invSimp ~ RestorationCategory + TotalFlowers + SemiNatural +
    (1 | Visit) + (1 | EasementID)
Data: bombus_responses

      AIC       BIC    logLik -2*log(L)  df.resid
    292.9     313.4    -138.4     276.9        88

Random effects:

Conditional model:
 Groups     Name        Variance Std.Dev.
 Visit      (Intercept) 0.130561 0.36133
 EasementID (Intercept) 0.005508 0.07421
 Residual               0.989584 0.99478
Number of obs: 96, groups:  Visit, 3; EasementID, 32

Dispersion estimate for gaussian family (sigma^2): 0.99

Conditional model:
                       Estimate Std. Error z value Pr(>|z|)
(Intercept)            0.603258   0.481127   1.254   0.2099
RestorationCategoryLow 0.543596   0.324353   1.676   0.0937 .
RestorationCategoryMod 0.827867   0.328347   2.521   0.0117 *
TotalFlowers           0.004888   0.001745   2.801   0.0051 **
SemiNatural            0.001719   0.006338   0.271   0.7862
---
Signif. codes:  0 '***' 0.001 '**' 0.01 '*' 0.05 '.' 0.1 ' ' 1
car::Anova(m2d)
Analysis of Deviance Table (Type II Wald chisquare tests)

Response: invSimp
                     Chisq Df Pr(>Chisq)
RestorationCategory 6.4243  2   0.040270 *
TotalFlowers        7.8447  1   0.005097 **
SemiNatural         0.0736  1   0.786181
---
Signif. codes:  0 '***' 0.001 '**' 0.01 '*' 0.05 '.' 0.1 ' ' 1
# pairwise comparisons
# uses TukeyHSD for multiple comparisons
emmeans(m2d, pairwise ~ RestorationCategory)
$emmeans
 RestorationCategory emmean    SE df lower.CL upper.CL
 No                    1.06 0.351 88    0.366     1.76
 Low                   1.61 0.269 88    1.072     2.14
 Mod                   1.89 0.263 88    1.368     2.41

Confidence level used: 0.95

$contrasts
 contrast  estimate    SE df t.ratio p.value
 No - Low    -0.544 0.324 88  -1.676  0.2201
 No - Mod    -0.828 0.328 88  -2.521  0.0356
 Low - Mod   -0.284 0.238 88  -1.195  0.4592

P value adjustment: tukey method for comparing a family of 3 estimates
# standardized estimates and transform to response scale
model_parameters(m2d,
                 standardize = "basic")
Warning: Standardizing coefficients only works for fixed effects of the mixed
  model.
# Fixed Effects

Parameter                 | Std. Coef. |   SE |        95% CI |    z |     p
----------------------------------------------------------------------------
(Intercept)               |       0.00 | 0.00 | [ 0.00, 0.00] | 1.25 | 0.210
RestorationCategory [Low] |       0.22 | 0.13 | [-0.04, 0.49] | 1.68 | 0.094
RestorationCategory [Mod] |       0.35 | 0.14 | [ 0.08, 0.61] | 2.52 | 0.012
TotalFlowers              |       0.28 | 0.10 | [ 0.09, 0.48] | 2.80 | 0.005
SemiNatural               |       0.02 | 0.09 | [-0.15, 0.20] | 0.27 | 0.786

Uncertainty intervals (equal-tailed) and p-values (two-tailed) computed
  using a Wald z-distribution approximation.
```

##### Differences in bumble bee community composition between sites with different restoration and management history

###### PERMANOVA: Community composition

```
set.seed(123) #so permutations are reproducible

p_anova1 <- adonis2(community_matrix ~
                      RestorationCategory +
                      TotalFlowers +
                      SemiNatural +
                      Year,
                    data = community_env,
                    method="bray",
                    by="margin",
                    permutations=999,
                    model = "reduced")
#model = "reduced" determines the method of permuations.
#This permutes the residuals under a reduced model

# view results
p_anova1
Permutation test for adonis under reduced model
Marginal effects of terms
Permutation: free
Number of permutations: 999

adonis2(formula = community_matrix ~ RestorationCategory + TotalFlowers + SemiNatural + Year, data = community_env, permutations = 999, method = "bray", by = "margin", model = "reduced")
                    Df SumOfSqs      R2      F Pr(>F)
RestorationCategory  2   0.5745 0.12199 2.3610  0.009 **
TotalFlowers         1   0.0934 0.01984 0.7679  0.574
SemiNatural          1   0.0475 0.01008 0.3902  0.858
Year                 1   0.0772 0.01638 0.6342  0.694
Residual            31   3.7717 0.80088
Total               36   4.7095 1.00000
---
Signif. codes:  0 '***' 0.001 '**' 0.01 '*' 0.05 '.' 0.1 ' ' 1
# pairwise permanova comparisons between restoration categories
pairwise.adonis(
  community_matrix,
  community_env$RestorationCategory,
  sim.function = "vegdist",
  sim.method = "bray")
       pairs Df SumsOfSqs  F.Model         R2 p.value p.adjusted sig
1 Mod vs Low  1 0.1545357 1.670059 0.05445241   0.131      0.393
2  Mod vs No  1 0.5767683 4.695776 0.19014490   0.004      0.012   .
3  Low vs No  1 0.3938919 2.592344 0.12005846   0.030      0.090
# Bray-Curtis dissimilarity matrix
bray1 <- vegdist(community_matrix, method = "bray")

# test for model assumption of dispersion
dispersion <- betadisper(
  bray1,
  group = community_env$RestorationCategory
  )

# view
dispersion

    Homogeneity of multivariate dispersions

Call: betadisper(d = bray1, group = community_env$RestorationCategory)

No. of Positive Eigenvalues: 16
No. of Negative Eigenvalues: 19

Average distance to median:
    No    Low    Mod
0.4456 0.2797 0.2538

Eigenvalues for PCoA axes:
(Showing 8 of 35 eigenvalues)
 PCoA1  PCoA2  PCoA3  PCoA4  PCoA5  PCoA6  PCoA7  PCoA8
1.5307 1.0472 0.6434 0.5779 0.4972 0.3889 0.3815 0.1770
permutest(dispersion, pairwise = T)

Permutation test for homogeneity of multivariate dispersions
Permutation: free
Number of permutations: 999

Response: Distances
          Df  Sum Sq  Mean Sq      F N.Perm Pr(>F)
Groups     2 0.16668 0.083340 4.1688    999  0.019 *
Residuals 34 0.67970 0.019991
---
Signif. codes:  0 '***' 0.001 '**' 0.01 '*' 0.05 '.' 0.1 ' ' 1

Pairwise comparisons:
(Observed p-value below diagonal, permuted p-value above diagonal)
          No      Low   Mod
No           0.064000 0.004
Low 0.063394          0.598
Mod 0.002287 0.595232
# no dispersion
```

### Figures

#### Bumble Abundance

##### Fig. 1A - restoration intervention intensity categories

```
# get predicted mean value for categories
# predict_response predicts for actual observed predictors
# this gives predictions on the responses scale
# use bias correction for back-transforming the predicted values
m3a_pr <- predict_response(m3a,
                           terms = c("RestorationCategory"),
                           interval = "confidence",
                           ci_level = 0.95,
                           margin = "marginalmeans",
                           type = "fixed",
                           bias_correction = TRUE)


# categorical plot to show mean effects of restoration
m3a_plot1 <- ggplot(m3a_pr) +
  geom_point(
    aes(x=x, y=predicted, fill=x, color=x),
    size = 6) +
  geom_linerange(
    aes(x = x, ymin = conf.low, ymax = conf.high, color=x),
    linewidth = 1.5) +
  labs(y = "Bumble Bee Abundance",
       x = "Restoration Intensity") +
  scale_color_manual(
    values = c("#767171", "#A9D18E", "#548235")) +
  theme_bw() +
  theme(
        axis.title = element_text(size = 18, face = "bold"),
        axis.text = element_text(size=12),
        axis.text.x = element_text(size = 16, face = "bold"),
        legend.position = "none",
        panel.grid.major = element_blank(),
        panel.grid.minor = element_blank(),
        plot.margin = margin(t = 15, r = 5, b = 5, l = 5)
        )

# view the plot
m3a_plot1
```

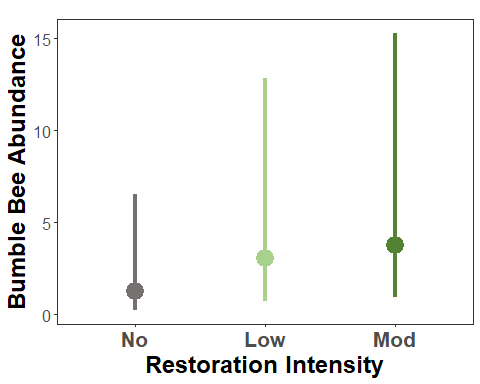

##### Fig. S3A - floral abundance

```
# get predicted mean value for categories
# predict_response predicts for actual observed predictors

# restrict to biologically realistic predictions
# dont want to extrapolate beyond data range
#summary(bombus_responses$TotalFlowers)
#   Min. 1st Qu.  Median    Mean 3rd Qu.    Max.
#   1.00   24.25   64.00   79.28  116.25  382.00

# Use observed data quantiles to set prediction range
max_flower <- quantile(bombus_responses$TotalFlowers, 0.99)
# 250

m3a_pr2 <- predict_response(m3a,
                            # restric the range here
                           terms = c("TotalFlowers [0:250]"),
                           interval = "confidence",
                           ci_level = 0.95,
                           margin = "marginalmeans",
                           type = "fixed",
                           bias_correction = TRUE)


# get the residuals for the predicted values
m3a_resid <- residualize_over_grid(m3a_pr2,
                                 m3a,
                                 protect_names = TRUE)


# continuous plot to show relationship between bees and flowers
# separated to show mean effects of restoration
m3a_plot2 <- ggplot(m3a_pr2) +
  geom_ribbon(aes(
    x = x,
    ymin = conf.low,
    ymax = conf.high),
    alpha = 0.1) +
  geom_point(data=m3a_resid %>%
               filter(!predicted > 200),
             aes(x=x,
                 y=predicted),
             color = "#bc87b2",
             size=4,
             alpha = 0.6) +
  geom_line(
    aes(x=x, y = predicted),
    color = "#775188",
    linewidth = 2) +

  labs(y = "Bumble Bee Abundance",
       x = "Floral Abundance") +
  theme_bw() +
  theme(
        axis.title = element_text(size = 18, face = "bold"),
        axis.text = element_text(size=12),
        axis.text.x = element_text(size = 16, face = "bold"),
        legend.position = "none",
        panel.grid.major = element_blank(),
        panel.grid.minor = element_blank(),
        plot.margin = margin(t = 15, r = 5, b = 5, l = 5)
        ) +
  scale_y_sqrt(breaks = c(0,5, 20, 50,100, 150),
                     limits = c(0,150))


m3a_plot2
```

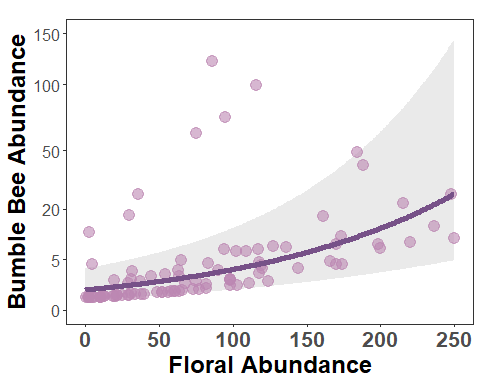

\# copy to clipboard 360 wide x 350 tall

##### Fig. S3C - landscape

```
# calculate predicted values
m3a_snh_pr <- predict_response(
  m3a,
  terms = c("SemiNatural [0:80]"),
  interval = "confidence",
  ci_level = 0.95,
  margin = "marginalmeans",
  type = "fixed",
  bias_correction = T)


# get the residuals for the predicted values
m3a_snh_resid <- residualize_over_grid(m3a_snh_pr,
                                 m3a,
                                 protect_names = TRUE)


# continuous plot to show relationship between bees and SNH
ggplot(m3a_snh_pr) +
  geom_ribbon(aes(
    x = x,
    ymin = (conf.low),
    ymax = (conf.high)),
    alpha = 0.1) +
   geom_point(data=m3a_snh_resid %>%
               filter(!predicted > 200),
             aes(x=x,
                 y=predicted),
             color = "#bc87b2",
             fill = "#bc87b2",
             size=4,
             alpha = 0.6,
             shape = 23) +
  geom_line(aes(
    x=x,
    y = (predicted)),
    color = "#775188",
    linewidth = 2,
    linetype = "dashed") +

  theme_bw() +
  theme(axis.title = element_blank(),
        #axis.title = element_text(size=16, face="bold"),
        axis.text = element_text(size=12),
        axis.text.x = element_text(size = 16, face = "bold"),
        panel.grid.major = element_blank(),
        panel.grid.minor = element_blank(),
        legend.position = "none"
        ) +
  labs(x = "% Semi Natural Habitat\n(in 1.5-km)",
       y = "Bumble Bee Abundance") +
  scale_y_sqrt(breaks = c(0,5, 20, 50,100, 150),
                     limits = c(0,150))
```

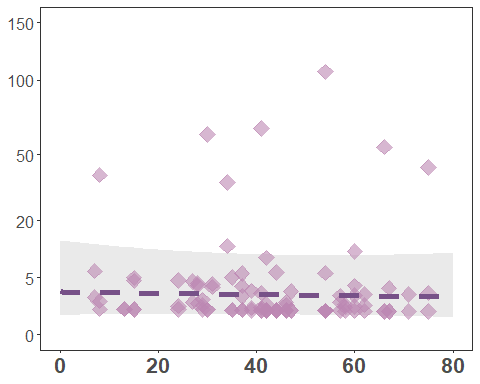

#### Diversity

##### Fig. 1B - restoration intervention intensity categories

```
# get predicted mean value for categories
# predict_response predicts for actual observed predictors
m2d_pr <- predict_response(m2d,
                           terms = c("RestorationCategory"),
                           interval = "confidence",
                           ci_level = 0.95,
                           margin = "marginalmeans",
                           type = "fixed",
                           bias_correction = TRUE)


# Create manual letter annotations for significance groups
sig_letters_m3d <- data.frame(
  x = c("No", "Low", "Mod"),
  label = c("a", "ab", "b"),  # example letters from Tukey HSD
  y = m2d_pr$conf.high + 0.15  # position letters slightly above upper CI
)


# categorical plot to show mean effects of restoration
m2d_plot <- ggplot(m2d_pr) +
  geom_point(
    aes(x=x, y=predicted, fill=x, color=x),
    size = 6) +
  geom_linerange(
    aes(x = x, ymin = conf.low, ymax = conf.high, color=x),
    linewidth = 1.5) +
  labs(y = "Bumble Bee Diversity",
       x = "Restoration Intensity") +
  # add significance annotations
  geom_text(
    data = sig_letters_m3d,
    aes(x = x, y = y, label = label),
    size = 6,
    fontface = "bold") +
  scale_color_manual(
    values = c("#767171", "#A9D18E", "#548235")) +
  theme_bw() +
  theme(
        axis.title = element_text(size = 18, face = "bold"),
        axis.text = element_text(size=12),
        axis.text.x = element_text(size = 16, face = "bold"),
        legend.position = "none",
        panel.grid.major = element_blank(),
        panel.grid.minor = element_blank(),
        plot.margin = margin(t = 15, r = 5, b = 5, l = 5)
        )

# view the plot
m2d_plot
```

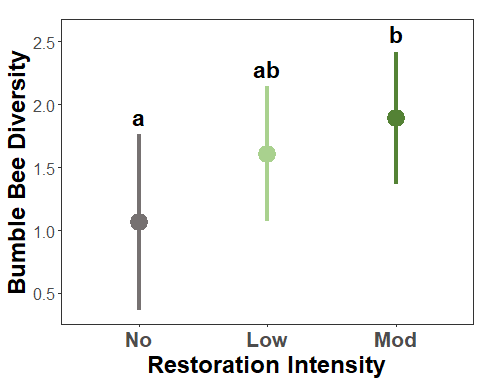

##### Fig. S3B - floral abundance

```
# get predicted mean value for categories
# predict_response predicts for actual observed predictors

# restrict to biologically realistic predictions
# dont want to extrapolate beyond data range
#summary(bombus_responses$TotalFlowers)
#   Min. 1st Qu.  Median    Mean 3rd Qu.    Max.
#   1.00   24.25   64.00   79.28  116.25  382.00

# Use observed data quantiles to set prediction range
max_flower <- quantile(bombus_responses$TotalFlowers, 0.99)
# 250

m2d_pr2 <- predict_response(m2d,
                           terms = c("TotalFlowers [0:250]"),
                           interval = "confidence",
                           ci_level = 0.95,
                           margin = "marginalmeans",
                           type = "fixed",
                           bias_correction = TRUE)


# dont want to extrapolate beyond data range
#summary(bombus_responses$TotalFlowers)
#   Min. 1st Qu.  Median    Mean 3rd Qu.    Max.
#   1.00   24.25   64.00   79.28  116.25  382.00

# get the residuals for the predicted values
m2d_resid <- residualize_over_grid(m2d_pr2,
                                 m2d,
                                 protect_names = TRUE)

# continuous plot to show relationship between bees and flowers
# separated to show mean effects of restoration
m2d_plot2 <- ggplot(m2d_pr2) +
  geom_ribbon(aes(
    x = x,
    ymin = conf.low,
    ymax = conf.high),
    alpha = 0.1) +
  geom_point(data=m2d_resid,
             aes(x=x,
                 y=predicted),
             color = "#8a9cbf",
             size=4,
             alpha = 0.6) +
  geom_line(
    aes(x=x,
        y = predicted),
    color = "#3d5591",
    linewidth = 2) +

  labs(y = "Bumble Bee Diversity",
       x = "Floral Abundance") +
  theme_bw() +
  theme(
        axis.title = element_text(size = 18, face = "bold"),
        axis.text = element_text(size=12),
        axis.text.x = element_text(size = 16, face = "bold"),
        legend.position = "none",
        panel.grid.major = element_blank(),
        panel.grid.minor = element_blank(),
        plot.margin = margin(t = 15, r = 5, b = 5, l = 5)
        ) +
  scale_y_continuous(breaks = c(0,1,2,3,4),
                     limits = c(-0.5,4.3))


# view the plot
m2d_plot2
```

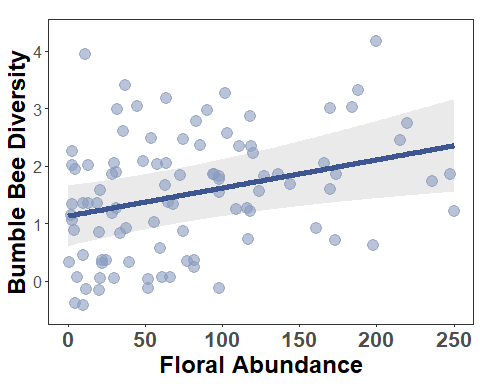

##### Fig. S3D - landscape

```
# calculate predicted values
m2d_snh_pr <- predict_response(
  m2d,
  terms = c("SemiNatural [0:80]"),
  interval = "confidence",
  ci_level = 0.95,
  margin = "marginalmeans",
  type = "fixed",
  bias_correction = T)


# get the residuals for the predicted values
m2d_snh_resid <- residualize_over_grid(m2d_snh_pr,
                                 m2d,
                                 protect_names = TRUE)


# continuous plot to show relationship between bees and SNH
ggplot(m2d_snh_pr) +
  geom_ribbon(aes(
    x = x,
    ymin = (conf.low),
    ymax = (conf.high)),
    alpha = 0.1) +
   geom_point(data=m2d_snh_resid,
             aes(x=x,
                 y=predicted),
             color = "#8a9cbf",
             fill = "#8a9cbf",
             size=4,
             alpha = 0.6,
             shape = 23) +
  geom_line(aes(
    x=x,
    y = (predicted)),
    color = "#3d5591",
    linewidth = 2,
    linetype = "dashed") +

  theme_bw() +
  theme(axis.title = element_blank(),
        #axis.title = element_text(size=16, face="bold"),
        axis.text = element_text(size=12),
        axis.text.x = element_text(size = 16, face = "bold"),
        panel.grid.major = element_blank(),
        panel.grid.minor = element_blank(),
        legend.position = "none"
        ) +
  labs(x = "% Semi Natural Habitat\n(in 1.5-km)",
       y = "Bumble Bee Diversity") +
  scale_y_continuous(breaks = c(0,1,2,3,4),
                     limits = c(-0.5,4.3))
```

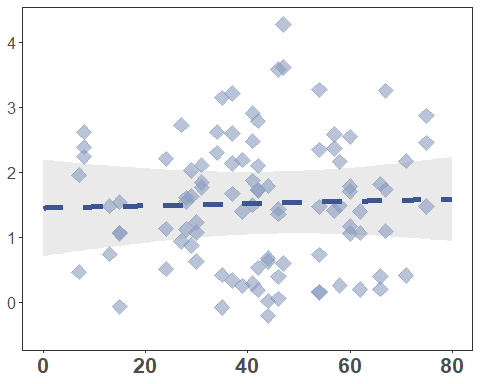

#### Fig. 2 - Community composition

```
# create the nmds with transformed data
nmds <- metaMDS(community_matrix, distance = "bray")
Run 0 stress 0.1276811
Run 1 stress 0.1723102
Run 2 stress 0.1277004
... Procrustes: rmse 0.004644083  max resid 0.02017576
Run 3 stress 0.1278812
... Procrustes: rmse 0.0198653  max resid 0.0956508
Run 4 stress 0.1276811
... Procrustes: rmse 5.051583e-05  max resid 0.000225221
... Similar to previous best
Run 5 stress 0.1276813
... Procrustes: rmse 0.001938201  max resid 0.008969975
... Similar to previous best
Run 6 stress 0.179262
Run 7 stress 0.1469172
Run 8 stress 0.1276813
... Procrustes: rmse 0.001940008  max resid 0.008981131
... Similar to previous best
Run 9 stress 0.1279005
... Procrustes: rmse 0.02004083  max resid 0.09583165
Run 10 stress 0.1277661
... Procrustes: rmse 0.009603677  max resid 0.04390416
Run 11 stress 0.1278811
... Procrustes: rmse 0.01986558  max resid 0.09564674
Run 12 stress 0.1468796
Run 13 stress 0.1354442
Run 14 stress 0.1276817
... Procrustes: rmse 0.002082865  max resid 0.009638706
... Similar to previous best
Run 15 stress 0.1276818
... Procrustes: rmse 0.002107036  max resid 0.009735828
... Similar to previous best
Run 16 stress 0.127708
... Procrustes: rmse 0.005692167  max resid 0.02516497
Run 17 stress 0.128263
Run 18 stress 0.1276813
... Procrustes: rmse 0.001858089  max resid 0.008590172
... Similar to previous best
Run 19 stress 0.1278804
... Procrustes: rmse 0.01987217  max resid 0.09563132
Run 20 stress 0.1387744
*** Best solution repeated 6 times
plot(nmds)

# remove site name column
community_envfit <- community_env[-c(1)]

(fit <- envfit(nmds, community_envfit, perm = 999))

***VECTORS

                NMDS1    NMDS2     r2 Pr(>r)
Year         -0.10794  0.99416 0.0270  0.691
SemiNatural   0.13674  0.99061 0.0145  0.797
TotalFlowers -0.74258 -0.66976 0.0697  0.300
Permutation: free
Number of permutations: 999

***FACTORS:

Centroids:
                         NMDS1   NMDS2
RestorationCategoryNo   0.6650  0.0438
RestorationCategoryLow -0.0804  0.0763
RestorationCategoryMod -0.1740 -0.0880

Goodness of fit:
                        r2 Pr(>r)
RestorationCategory 0.1794  0.009 **
---
Signif. codes:  0 '***' 0.001 '**' 0.01 '*' 0.05 '.' 0.1 ' ' 1
Permutation: free
Number of permutations: 999
head(fit)
$vectors
                NMDS1    NMDS2     r2 Pr(>r)
Year         -0.10794  0.99416 0.0270  0.691
SemiNatural   0.13674  0.99061 0.0145  0.797
TotalFlowers -0.74258 -0.66976 0.0697  0.300
Permutation: free
Number of permutations: 999

$factors
Centroids:
                         NMDS1   NMDS2
RestorationCategoryNo   0.6650  0.0438
RestorationCategoryLow -0.0804  0.0763
RestorationCategoryMod -0.1740 -0.0880

Goodness of fit:
                        r2 Pr(>r)
RestorationCategory 0.1794  0.009 **
---
Signif. codes:  0 '***' 0.001 '**' 0.01 '*' 0.05 '.' 0.1 ' ' 1
Permutation: free
Number of permutations: 999

$na.action
function (object, ...)
UseMethod("na.action")
<bytecode: 0x00000184f7b932d8>
<environment: namespace:stats>
# get score values for the grouping vectors
scores(fit, "vectors")
                   NMDS1      NMDS2
Year         -0.01775148  0.1634939
SemiNatural   0.01649079  0.1194673
TotalFlowers -0.19607581 -0.1768464
# create a data frame with the scores
env_scores <- as.data.frame(scores(fit, display = "vectors"))
env_scores <- cbind(env_scores, Species = rownames(env_scores))

# view the base plot for overall pattern
plot(nmds)
plot(fit, col="black")
```

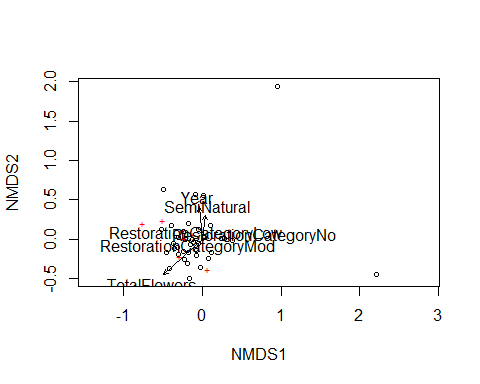

```
# extract NMDS scores (x and y coordinates)
# need scores in a dateframe in order to make a nmds plot using ggplot
# then add identifying columns to dataframe
# ggplot needs these in a df, so specify tidy = TRUE
nmds_sites <- scores(nmds, display = c("sites", "species"), tidy = TRUE)

nmds_scores <- vegan::scores(nmds, display = "sites") %>%
  as.data.frame() %>%
  bind_cols(community_env) %>%
  mutate(
    Group = factor(RestorationCategory,
                   levels = c("No", "Low", "Mod")))


# basic hull
# create hulls
grp_no <- nmds_scores[nmds_scores$RestorationCategory == "No", ][chull(nmds_scores[nmds_scores$RestorationCategory ==
    "No", c("NMDS1", "NMDS2")]), ]  # hull values for grp A

grp_low <- nmds_scores[nmds_scores$RestorationCategory == "Low", ][chull(nmds_scores[nmds_scores$RestorationCategory ==
    "Low", c("NMDS1", "NMDS2")]), ]  # hull values for grp B

grp_mod <- nmds_scores[nmds_scores$RestorationCategory == "Mod", ][chull(nmds_scores[nmds_scores$RestorationCategory ==
    "Mod", c("NMDS1", "NMDS2")]), ]  # hull values for grp C

#combine groups' data
hull_data <- rbind(grp_no, grp_low, grp_mod) %>%
  mutate(
    Group = factor(RestorationCategory,
                   levels = c("No", "Low", "Mod")))


# make the final plot
plot_nmds <- ggplot() +
  geom_polygon(data=hull_data,
               aes(x=NMDS1,y=NMDS2, group=Group,
                   linetype = Group, fill = Group),
               alpha=0.2, color="black") +

  geom_point(data = nmds_scores,
             aes(x=NMDS1,y=NMDS2,
                 shape = Group, fill = Group),
             size = 5) + # add the point markers

  # create custom and consistent color and line mappings
  scale_fill_manual(
    values = c("No" = "#767171", "Low" ="#A9D18E","Mod" ="#548235"),
    name = "Restoration\nIntensity",
    guide = "legend"
                    ) +
  scale_color_manual(
    values = c("No" = "#767171", "Low" = "#A9D18E", "Mod"="#548235"),
    name = "Restoration\nIntensity",
    guide = "legend") +

  scale_shape_manual(
    values = c("No" =21,"Low" =22,"Mod" =24),
    name = "Restoration\nIntensity",
    guide = "legend") +

  scale_linetype_manual(
    values = c("No"="solid", "Low"="dashed", "Mod"="longdash"),
    name = "Restoration\nIntensity",
    guide = "legend") +

  # customize the plot theme
  theme_bw() +
  theme(axis.text.x = element_blank(),  # remove x-axis text
        axis.text.y = element_blank(), # remove y-axis text
        axis.ticks = element_blank(),  # remove axis ticks
        axis.title = element_text(size=16, face = "bold"),
        panel.background = element_blank(),
        panel.grid.major = element_blank(),  #remove major-grid labels
        panel.grid.minor = element_blank(),  #remove minor-grid labels
        plot.background = element_blank(),
        legend.title = element_text(size=14),
        legend.background = element_rect(color = "black",linetype = "solid"),
        legend.text = element_text(size=16, face="bold"),
        legend.position = "right") +
  labs(x = "NMDS1", y = "NMDS2")

plot_nmds
Warning: No shared levels found between `names(values)` of the manual scale and the
data's colour values.
```

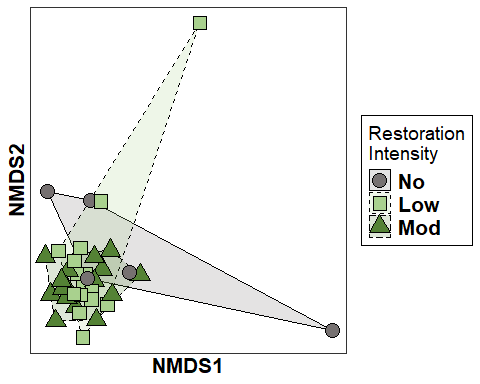

```
# extra information
nmds$stress
[1] 0.1276811
stressplot(nmds)
```

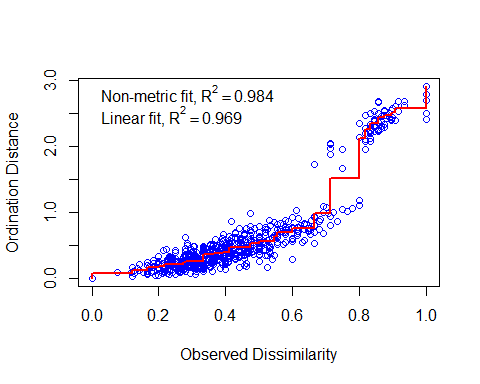

#### Floral Abundance

##### Fig. 3 - restoration intervention intensity categories

```
# get predicted mean value for categories
# predict_response predicts for actual observed predictors
mfa_pr <- predict_response(m1fa,
                           terms = c("RestorationCategory"),
                           interval = "confidence",
                           ci_level = 0.95,
                           margin = "marginalmeans",
                           type = "fixed",
                           bias_correction = T)
NOTE: Results may be misleading due to involvement in interactions
# Create manual letter annotations for significance groups
sig_letters_mfa <- data.frame(
  x = c("No", "Low", "Mod"),
  label = c("a", "b", "c"),  # example letters from Tukey HSD
  y = mfa_pr$conf.high + 10  # position letters slightly above upper CI
)


# categorical plot to show mean effects of restoration
mfa_plot1 <- ggplot(mfa_pr) +
  geom_point(aes(
    x=x,
    y=predicted,
    fill=x,
    color=x),
    size = 6) +
  geom_linerange(aes(
    x=x,
    ymin=conf.low,
    ymax=conf.high,
    color=x),
    linewidth = 1.5) +
  labs(y = "Floral Abundance",
       x = "Restoration Intensity") +

  scale_color_manual(
    values = c("#767171", "#A9D18E", "#548235")) +

#   add significance annotations
  geom_text(data = sig_letters_mfa,
            aes(x = x, y = y, label = label),
            size = 6, fontface = "bold") +

  theme_bw() +
  theme(axis.title = element_text(size=18, face="bold"),
        axis.text = element_text(size=12),
        axis.text.x = element_text(size = 16, face = "bold"),
        legend.position = "none",
        panel.grid.major = element_blank(),
        panel.grid.minor = element_blank())

# view the plot
mfa_plot1
```

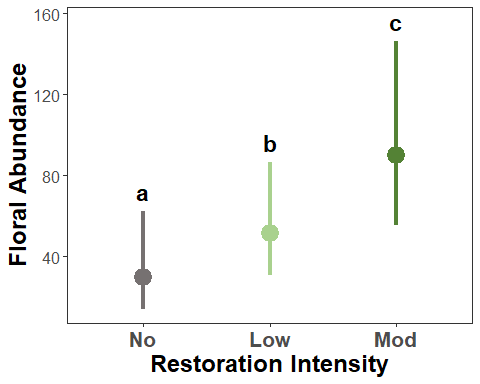

\# copy to clipboard 360 wide x 350 tall

##### Fig. S5 - landscape

```
# continuous plot to show relationship between bees and flowers
# separated to show mean effects of restoration

#summary(bombus_responses$SemiNatural)
#   Min. 1st Qu.  Median    Mean 3rd Qu.    Max.
#  6.992  30.461  42.224  42.222  57.492  75.452

mfa_pr2 <- predict_response(m1fa,
                           terms = c("SemiNatural [0:80]", #restrict
                                     "RestorationCategory"),
                           interval = "confidence",
                           ci_level = 0.95,
                           margin = "marginalmeans",
                           type = "fixed",
                           bias_correction = T)


# get the residuals for the predicted values
mfa_resid <- residualize_over_grid(mfa_pr2,
                                 m1fa,
                                 protect_names = TRUE)


# continuous plot
# relationship between floral and SNH x restoration
ggplot(mfa_pr2) +
  geom_ribbon(aes(
    x = x,
    ymin = conf.low,
    ymax = conf.high,
    fill = group),
    alpha = 0.1) +
  geom_point(data=mfa_resid %>%
               filter(!predicted > 350),
             aes(x=x,
                 y=predicted,
                 color = group),
             size=4,
             alpha = 0.5) +
  geom_line(
    aes(x=x, y = predicted,
        color = group,
        linetype = group),
    linewidth = 2) +
  scale_color_manual(
    values = c("#767171", "#A9D18E", "#548235"),
    name = "Restoration\nIntensity",
                    guide = "legend") +
  scale_fill_manual(
    values = c("#767171", "#A9D18E", "#548235"),
    name = "Restoration\nIntensity",
                    guide = "legend") +
  scale_linetype_manual(
    values = c("No"="solid", "Low"="dashed", "Mod"="dashed"),
    name = "Restoration\nIntensity",
                    guide = "legend") +

  labs(y = "Floral Abundance",
       x = "% Semi Natural Habitat\n(in 1.5-km)") +
  theme_bw() +
  theme(axis.title = element_text(size=16, face="bold"),
        axis.text = element_text(size=12),
        axis.text.x = element_text(size = 16, face = "bold"),
        #legend.position = "none",
        panel.grid.major = element_blank(),
        panel.grid.minor = element_blank(),
       # plot.margin = margin(t = 15, r = 5, b = 5, l = 5),
        legend.title = element_text(size=14),
        legend.background = element_rect(color = "black",
                                         linetype = "solid"),
        legend.text = element_text(size=16, face="bold")
        )
```

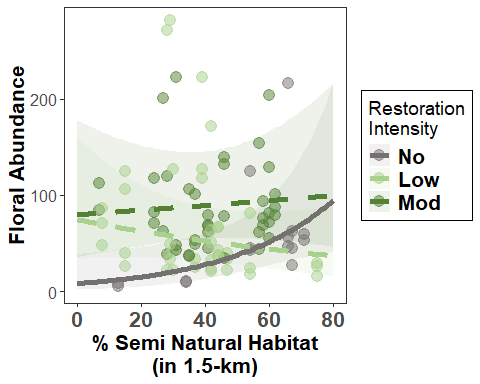

### Pairwise contrasts

#### Bumble abundance

```
# (1) generate estimated marginal means by
# restoration intervention intensity
# this is on the default link (log) scale
m3a_emm <- emmeans(m3a, ~ RestorationCategory)

#############################################
# compare no intervention to ANY intervention
#############################################

# (2) define custom contrast
# No intervention vs any intervention (Low + Mod)
# (0.5 * Low + 0.5 * Mod) - (1 * No)
m3a_contrast <- contrast(
  m3a_emm,
  # specify the (-) so positive results = increase with restoration
  method = list("Restoration vs No" = c(-1, 0.5, 0.5))
)

# (3) get confidence intervals
# (still log scale)
m3a_contrast_ci <- confint(m3a_contrast)

# (4) combine information and calculate to percent change
# finally convert to the response scale
m3a_contrast_summary <- as.data.frame(m3a_contrast_ci) %>%
  left_join(as.data.frame(m3a_contrast)
            [, c("contrast", "SE", "z.ratio", "p.value")],
            by = c("contrast", "SE")) %>%
  mutate(
    # exponentiate: get ratio of means (Restoration / No)
    ratio = exp(estimate),
    # convert to percent change
    percent.change = (ratio - 1) * 100,
    lower.CL = (exp(asymp.LCL) - 1) * 100,
    upper.CL = (exp(asymp.UCL) - 1) * 100
  ) %>%
  # round and select relevant columns
  mutate(across(
    c(ratio, percent.change, lower.CL, upper.CL), round, 1)) %>%
  select(contrast, ratio, percent.change, lower.CL, upper.CL,
         SE, z.ratio, p.value)
Warning: There was 1 warning in `mutate()`.
ℹ In argument: `across(c(ratio, percent.change, lower.CL, upper.CL), round,
  1)`.
Caused by warning:
! The `...` argument of `across()` is deprecated as of dplyr 1.1.0.
Supply arguments directly to `.fns` through an anonymous function instead.

  # Previously
  across(a:b, mean, na.rm = TRUE)

  # Now
  across(a:b, \(x) mean(x, na.rm = TRUE))
m3a_contrast_summary
           contrast ratio percent.change lower.CL upper.CL        SE  z.ratio
1 Restoration vs No   2.7          168.3     -4.2    651.5 0.5255418 1.877852
     p.value
1 0.06040139
#############################################
# compare all intervention intensities
#############################################

# (2) define custom contrast
# gives Low - No, Mod - No, Mod - Low
p2 <- pairs(m3a_emm, reverse = TRUE)

# (3) get confidence intervals
# (still log scale)
p_ci2 <- confint(p2)

# (4) combine information and calculate to percent change
# finally convert to the response scale
m3a_summary2 <- as.data.frame(p_ci2) %>%
  left_join(as.data.frame(p2)
            [, c("contrast", "SE", "z.ratio", "p.value")],
            by = c("contrast", "SE")) %>%
  mutate(
    # exponentiate: get ratio of means (Restoration / No)
    ratio = exp(estimate),
    # convert to the relative difference & percent change
    percent.change = (ratio - 1) * 100,
    lower.CL = (exp(asymp.LCL) - 1) * 100,
    upper.CL = (exp(asymp.UCL) - 1) * 100
  ) %>%
  # round and select relevant columns
  mutate(across(
    c(ratio, percent.change, lower.CL, upper.CL), round, 1)) %>%
  select(
    contrast, ratio, percent.change, lower.CL, upper.CL,
    SE, z.ratio, p.value)

m3a_summary2
   contrast ratio percent.change lower.CL upper.CL        SE   z.ratio
1  Low - No   2.4          143.0    -32.9    780.1 0.5491357 1.6168787
2  Mod - No   3.0          196.2    -18.8    980.7 0.5522596 1.9662739
3 Mod - Low   1.2           21.9    -43.6    163.6 0.3290969 0.6016704
    p.value
1 0.2384401
2 0.1206325
3 0.8191951
```

#### Diversity

```
# (1) generate estimated marginal means by
# restoration intervention intensity
# this is on the default link (log) scale
m2d_emm <- emmeans(m2d, ~ RestorationCategory)

#############################################
# compare no intervention to ANY intervention
#############################################

# (2) define custom contrast
# No intervention vs any intervention (Low + Mod)
# (0.5 * Low + 0.5 * Mod) - (1 * No)
m2d_contrast <- contrast(
  m2d_emm,
  # specify the (-) so positive results = increase with restoration
  method = list("Restoration vs No" = c(-1, 0.5, 0.5))
)

# (3) get confidence intervals
# (still log scale)
m2d_contrast_ci <- confint(m2d_contrast)

# (4) combine information and calculate to percent change
# finally convert to the response scale
m2d_contrast_summary <- as.data.frame(m2d_contrast_ci) %>%
  left_join(as.data.frame(m3a_contrast)
            [, c("contrast", "SE", "z.ratio", "p.value")],
            by = c("contrast", "SE")) %>%
  mutate(
    # exponentiate: get ratio of means (Restoration / No)
    ratio = exp(estimate),
    # convert to percent change
    percent.change = (ratio - 1) * 100,
    lower.CL = (exp(lower.CL) - 1) * 100,
    upper.CL = (exp(upper.CL) - 1) * 100
  ) %>%
  # round and select relevant columns
  mutate(across(
    c(ratio, percent.change, lower.CL, upper.CL), round, 1)) %>%
  select(contrast, ratio, percent.change, lower.CL, upper.CL,
         SE, z.ratio, p.value)

m2d_contrast_summary
           contrast ratio percent.change lower.CL upper.CL        SE z.ratio
1 Restoration vs No     2           98.5      8.5    263.2 0.3039195      NA
  p.value
1      NA
#############################################
# compare all intervention intensities
#############################################

# (2) define custom contrast
# gives Low - No, Mod - No, Mod - Low
p2d <- pairs(m2d_emm, reverse = TRUE)

# (3) get confidence intervals
# (still log scale)
p2d_ci2 <- confint(p2d)

# (4) combine information and calculate to percent change
# finally convert to the response scale
m2d_summary2 <- as.data.frame(p2d_ci2) %>%
  left_join(as.data.frame(p2)
            [, c("contrast", "SE", "z.ratio", "p.value")],
            by = c("contrast", "SE")) %>%
  mutate(
    # exponentiate: get ratio of means (Restoration / No)
    ratio = exp(estimate),
    # convert to the relative difference & percent change
    percent.change = (ratio - 1) * 100,
    lower.CL = (exp(lower.CL) - 1) * 100,
    upper.CL = (exp(upper.CL) - 1) * 100
  ) %>%
  # round and select relevant columns
  mutate(across(
    c(ratio, percent.change, lower.CL, upper.CL), round, 1)) %>%
  select(
    contrast, ratio, percent.change, lower.CL, upper.CL,
    SE, z.ratio, p.value)

m2d_summary2
   contrast ratio percent.change lower.CL upper.CL        SE z.ratio p.value
1  Low - No   1.7           72.2    -20.5    273.2 0.3243527      NA      NA
2  Mod - No   2.3          128.8      4.6    400.6 0.3283471      NA      NA
3 Mod - Low   1.3           32.9    -24.6    134.3 0.2378332      NA      NA
```

#### Floral abundance

##### restoration intervention intensity

```
# (1) generate estimated marginal means by
# restoration intervention intensity
# this is on the default link (log) scale
mfa_emm <- emmeans(m1fa,
                   ~ RestorationCategory*SemiNatural)

#############################################
# compare all intervention intensities
#############################################

# (2) define custom contrast
# gives Low - No, Mod - No, Mod - Low
# this is for the median value of SNH (42.2)
pf <- pairs(mfa_emm,
             reverse = TRUE,
             by = "SemiNatural")

# (3) get confidence intervals
# (still log scale)
pf_ci <- confint(pf)

# (4) combine information and calculate to percent change
# finally convert to the response scale
mfa_summary <- as.data.frame(pf_ci) %>%
  left_join(as.data.frame(pf)
            [, c("contrast", "SE", "z.ratio", "p.value")],
            by = c("contrast", "SE")) %>%
  mutate(
    # exponentiate: get ratio of means (Restoration / No)
    ratio = exp(estimate),
    # convert to the relative difference & percent change
    percent.change = (ratio - 1) * 100,
    lower.CL = (exp(asymp.LCL) - 1) * 100,
    upper.CL = (exp(asymp.UCL) - 1) * 100
  ) %>%
  # round and select relevant columns
  mutate(across(
    c(ratio, percent.change, lower.CL, upper.CL), round, 1)) %>%
  select(
    contrast, ratio, percent.change, lower.CL, upper.CL,
    SE, z.ratio, p.value)

mfa_summary
   contrast ratio percent.change lower.CL upper.CL        SE  z.ratio
1  Low - No   1.7           73.5    -18.4    268.9 0.3217763 1.712968
2  Mod - No   3.0          201.6     44.7    528.7 0.3133820 3.522810
3 Mod - Low   1.7           73.8     13.1    167.1 0.1834046 3.014063
      p.value
1 0.200271168
2 0.001241624
3 0.007272119
```

##### landscape

```
############################################
# (1) pick representative SemiNatural values
############################################
sn_vals <- quantile(
  floral_incidence$SemiNatural,
  probs = c(0.1, 0.5, 0.9))

sn_vals
     10%      50%      90%
15.42872 42.22392 65.70227
##################################################
# (2) estimate marginal means at these SNH values
##################################################

mfa_emm_sn <- emmeans(m1fa,
                      ~ RestorationCategory*SemiNatural,
                      at = list(SemiNatural = sn_vals),
                      # response scale (back-transformed)
                      type = "response")


mfa_emm_sn_df <- as.data.frame(mfa_emm_sn) %>%
  select(RestorationCategory, SemiNatural, response, SE,
         asymp.LCL, asymp.UCL)

########################
# (3) contrasts
########################

# Pairwise for low vs. high SNH within each restoration category
mfa_snh_contrasts <- contrast(
  mfa_emm_sn,
  method = list("High vs Low" = c(-1, 0, 1)),  # compares 10th → 90th percentile
  by = "RestorationCategory",
  adjust = "fdr"  # adjust p-values for multiple comparisons
)


# extract and combine predicted means into a df
mfa_snh_contrasts_df <- as.data.frame(mfa_snh_contrasts) %>%
  select(RestorationCategory, contrast, ratio, SE, z.ratio, p.value) %>%
  mutate(
    percent.change = (ratio - 1) * 100
  )

###########################################################
# (4) combine information and calculate to percent change
##########################################################

mfa_snh_summary <- mfa_emm_sn_df %>%
  group_by(RestorationCategory) %>%
  summarize(
    SNH_low = min(SemiNatural),
    SNH_high = max(SemiNatural),
    Floral_low = response[SemiNatural == SNH_low],
    Floral_high = response[SemiNatural == SNH_high],
    Low_LCL = asymp.LCL[SemiNatural == SNH_low],
    Low_UCL = asymp.UCL[SemiNatural == SNH_low],
    High_LCL = asymp.LCL[SemiNatural == SNH_high],
    High_UCL = asymp.UCL[SemiNatural == SNH_high]
  ) %>%
  left_join(mfa_snh_contrasts_df %>%
              select(RestorationCategory, ratio, percent.change, p.value),
            by = "RestorationCategory") %>%
   # round and select relevant columns
  mutate(across(
    c(
      ratio, SNH_low, SNH_high, Floral_low, Floral_high,
      percent.change, Low_LCL, Low_UCL, High_LCL,High_UCL),
    round, 1),
    across(c(p.value), round, 3)
    ) %>%
  select(RestorationCategory, ratio, percent.change, p.value)


mfa_snh_summary
# A tibble: 3 × 4
  RestorationCategory ratio percent.change p.value
  <fct>               <dbl>          <dbl>   <dbl>
1 No                    4.6          363.    0.014
2 Low                   0.6          -35.6   0.281
3 Mod                   1.1           15     0.725
```
