## Supplementary Material for "Bumble Bee Abundance and Diversity Increase with Intensity of Tallgrass Prairie Restoration Intervention"

**Supporting Information**

**Supplementary Tables and Figures**

**Table S1.** Summary of fitted models testing the effects of restoration intervention intensity and semi-natural habitat in the surrounding landscape (SNH) on bumble bee and floral responses. The random effects of year, visit, and site were considered to account for interannual and seasonal variation and repeated measures. Results show the error structure for each model, the estimated coefficient (*β*), test statistic, and significance (*p*) for fixed effects. The overall variance explained by the fixed effects (*R^2^_m_*) and Akaike Information Criterion (AIC) are also shown. All GLMMs were systematically checked for data overdispersion, and residuals were checked for normality and heteroscedasticity using the *DHARMa* package (v0.4.7; Hartig, 2016). Significant predictors are indicated by asterisks: p < 0.05 (*), p < 0.01 (**), and p < 0.001 (***). The final models used for inference for each response variable are indicated in bold.

| **Model Formula** | **Error** | ***β*** | ***SE*** | ***p*** | ***R^2^_m_*** | **AIC** |
| --- | --- | --- | --- | --- | --- | --- |
| ***Bumble Bee Abundance*** |  |  |  |  |  |  |
| Restoration*SNH + Floral*SNH +  (1 \| Year) + (1 \| Visit) + (1\| Site) | poisson | - | - | - | 0.308 | 506.1 |
| Intercept | - | -2.10 | 1.66 | 0.207 | - | - |
| Restoration [Low] | - | 1.4 | 1.54 | 0.364 | - | - |
| Restoration [Mod] | - | 3.2 | 1.57 | 0.042* | - | - |
| SNH | - | 0.01 | 0.03 | 0.667 | - | - |
| Floral | - | 0.09 | 0.003 | 0.001** | - | - |
| Restoration [Low] x SNH | - | 0.06 | 0.03 | 0.844 | - | - |
| Restoration [Mod] x SNH | - | -0.04 | 0.03 | 0.204 | - | - |
| SNH x Floral | - | 9.4e-05 | 6.3e-05 | 0.136 | - | - |
| Restoration + Floral + SNH +  (1 \| Year) + (1 \| Visit) + (1\| Site) | poisson | - | - | - | 0.308 | 505.3 |
| Intercept | - | -1.76 | 1.17 | 0.132 | - | - |
| Restoration [Low] | - | 1.41 | 0.60 | 0.019* | - | - |
| Restoration [Mod] | - | 1.52 | 0.60 | 0.011* | - | - |
| Floral | - | 0.01 | 0.001 | <0.001*** | - | - |
| SNH | - | 5e-02 | 0.01 | 0.613 | - | - |
| Restoration + Floral + SNH +  (1 \| Year) + (1 \| Visit) + (1\| Site) | nbinom1 | - | - | - | 0.310 | 424.2 |
| Intercept | - | -0.25 | 0.89 | 0.783 | - | - |
| Restoration [Low] | - | 0.52 | 0.52 | 0.314 | - | - |
| Restoration [Mod] | - | 0.95 | 0.49 | 0.058 | - | - |
| Floral | - | 8.5e-03 | 0.002 | <0.001*** | - | - |
| SNH | - | 1.4e-05 | 0.01 | 0.999 | - | - |
| **Restoration + Floral + SNH +**  **(1 \| Year) + (1 \| Visit) + (1\| Site)** | **nbinom2** | - | - | - | **0.389** | **425.2** |
| **Intercept** | - | **-1.05** | **0.94** | **0.264** | - | - |
| **Restoration [Low]** | - | **0.89** | **0.55** | **0.1059** | - | - |
| **Restoration [Mod]** | - | **1.09** | **0.55** | **0.0493** | - | - |
| **Floral** | - | **0.01** | **0.003** | **<0.001***** | - | - |
| **SNH** | - | **-3e-03** | **0.01** | **0.7257** | - | - |
| Restoration + Floral +  (1 \| Year) + (1 \| Visit) + (1\| Site) | nbinom2 | - | - | - | 0.387 | 423.3 |
| Intercept | - | -1.21 | 0.83 | 0.145 | - | - |
| Restoration [Low] | - | 0.92 | 0.54 | 0.087 | - | - |
| Restoration [Mod] | - | 1.12 | 0.55 | 0.034* | - | - |
| Floral | - | 0.01 | 0.003 | <0.001*** | - | - |
| **Model Formula** | **Error** | ***β*** | *SE* | ***p*** | ***R^2^_m_*** | **AIC** |
| ***Bumble Bee Diversity*** |  |  |  |  |  |  |
| Restoration + Floral + SNH +  (1 \| Year) + (1 \| Visit) + (1\| Site) | gaussian | - | - | - | 0.202 | 294.9 |
| Intercept | - | 0.60 | 0.48 | 0.210 | - | - |
| Restoration [Low] | - | 0.54 | 0.32 | 0.094 | - | - |
| Restoration [Mod] | - | 0.83 | 0.33 | 0.012* | - | - |
| Floral | - | 0.005 | 0.002 | 0.005** | - | - |
| SNH | - | 0.002 | 0.01 | 0.786 | - | - |
| **Restoration + Floral + SNH +**  **(1 \| Visit) + (1\| Site)** | **gaussian** | - | - | - | **0.182** | **292.9** |
| **Intercept** | - | **0.60** | **0.48** | **0.210** | - | - |
| **Restoration [Low]** | - | **0.54** | **0.32** | **0.094** | - | - |
| **Restoration [Mod]** | - | **0.83** | **0.33** | **0.012*** | - | - |
| **Floral** | - | **0.005** | **0.002** | **0.005**** | - | - |
| **SNH** | - | **0.002** | **0.01** | **0.786** | - | - |
| Restoration + Floral +  (1 \| Visit) + (1\| Site) | gaussian | - | - | - | 0.182 | 290.9 |
| Intercept | - | 0.70 | 0.34 | 0.040 * | - | - |
| Restoration [Low] | - | 0.52 | 0.31 | 0.095 | - | - |
| Restoration [Mod] | - | 0.81 | 0.32 | 0.012* | - | - |
| Floral | - | 0.005 | 0.002 | 0.005** | - | - |
| **Model Formula** | **Error** | ***β*** | **SE** | ***p*** | ***r^2^*** | **AIC** |
| ***Floral Abundance*** |  |  |  |  |  |  |
| Restoration*SNH +  (1 \| Year) + (1 \| Visit) + (1\| Site) | poisson | - | - | - | - | 2786.3 |
| Intercept | - | 0.34 | 0.73 | 0.639 | - | - |
| Restoration [Low] | - | 4.15 | 0.77 | <0.001*** | - | - |
| Restoration [Mod] | - | 3.97 | 0.78 | <0.001*** | - | - |
| SNH | - | 0.05 | 0.01 | <0.001*** | - | - |
| Restoration [Low] x SNH | - | -0.07 | 0.02 | <0.001*** | - | - |
| Restoration [Mod] x SNH | - | -0.05 | 0.02 | <0.001*** | - | - |
| **Restoration*SNH +**  **(1 \| Visit) + (1\| Site)** | **nbinom1** | - |  | - | **0.403** | **1029.7** |
| **Intercept** | - | **2.09** | **0.79** | **0.008 **** | - | - |
| **Restoration [Low]** | - | **2.20** | **0.81** | **0.007**** | - | - |
| **Restoration [Mod]** | - | **2.27** | **0.82** | **0.006**** | - | - |
| **SNH** | - | **0.03** | **0.01** | **0.014*** | - | - |
| **Restoration [Low] x SNH** | - | **-0.04** | **0.02** | **0.008**** | - | - |
| **Restoration [Mod] x SNH** | - | **-0.03** | **0.02** | **0.061** | - | - |
| Restoration*SNH +  (1 \| Visit) + (1\| Site) | nbinom2 | - | - | - | 0.306 | 1028.2 |
| Intercept | - | 0.61 | 0.82 | 0.454 | - | - |
| Restoration [Low] | - | 4.04 | 0.87 | <0.001*** | - | - |
| Restoration [Mod] | - | 3.74 | 0.88 | <0.001*** | - | - |
| SNH | - | 0.05 | 0.01 | <0.001*** | - | - |
| Restoration [Low] x SNH | - | -0.07 | 0.02 | <0.001*** | - | - |
| Restoration [Mod] x SNH | - | -0.05 | 0.02 | 0.002** | - | - |

**Table S2.** List of bumble bee (*Bombus* spp.) species and total number of individuals observed for each, ordered from most to least abundant across sites in each type of restoration: no intervention (*n* = 6), low intensity intervention (*n* = 13), moderate intensity (*n* = 13). * Species of conservation concern; †not used in community NMDS analysis.

| **Latin species name** | **Common name** | **No** | **Low** | **Mod** |
| --- | --- | --- | --- | --- |
| *impatiens* | Common Eastern | 22 | 310 | 313 |
| *griseocollis* | Brown-Belted | 36 | 135 | 380 |
| *auricomus / pensylvanicus* | Black and Gold / American | 1 | 39 | 94 |
| *vagans / sandersoni* | Half-Black / Sanderson’s | 7 | 25 | 48 |
| *rufocinctus* | Red-Belted | 4 | 23 | 43 |
| *bimaculatus* | Two-Spotted | 1 | 27 | 37 |
| *borealis* † | Northern Amber | 1 | 4 | 15 |
| *affinis* *† | Rusty-Patched | 0 | 4 | 8 |
| *fervidus* *† | Golden Northern | 0 | 4 | 7 |
| *citrinus* † | Lemon Cuckoo | 0 | 3 | 0 |

**Table S3.** Flowering plant species and total number of occurrences of each, sorted in descending order of abundance across all sites in each type of restoration: no intervention (“No”, *n* = 6), low intensity intervention (“Low”, *n* = 13), moderate intensity (“Mod”, *n* = 13). For simplicity, species shown represent ≥ 80% of total plant occurrences. Species used by *Bombus affinis* based on Wolf et al. 2022 and personal observations are denoted with (*) and (**), respectively.

| **Latin binomial name** | **Common name** | **No** | **Low** | **Mod** |
| --- | --- | --- | --- | --- |
| *Monarda fistulosa ** | Bee balm / Wild bergamot | 0 | 449 | 854 |
| *Ratibida pinnata ** | Gray coneflower | 0 | 147 | 508 |
| *Rudbeckia hirta ** | Black-eyed Susan | 0 | 171 | 353 |
| *Erigeron strigosus* | Daisy or rough fleabane | 21 | 205 | 158 |
| *Desmodium canadense ** | Canada tick-trefoil | 1 | 7 | 343 |
| *Trifolium hybridum *** | Alsike clover | 3 | 176 | 84 |
| *Solidago canadensis ** | Canada goldenrod | 3 | 65 | 132 |
| *Heliopsis helianthoides* | False sunflower | 0 | 0 | 172 |
| *Medicago lupulina* | Black medick | 6 | 30 | 134 |
| *Silphium perfoliatum ** | Cup-plant | 0 | 71 | 90 |
| *Ambrosia trifida* | Giant ragweed | 34 | 93 | 33 |
| *Lotus corniculatus ** | Bird’s-foot trefoil | 136 | 3 | 1 |
| *Persicaria maculosa* | Spotted lady’s thumb | 0 | 111 | 0 |
| *Helianthus hirsutus* | Hairy or rough sunflower | 0 | 0 | 110 |
| *Penstemon digitalis* | Foxglove beardtongue | 0 | 15 | 86 |
| *Cirsium arvense* | Canada thistle | 1 | 60 | 39 |
| *Berteroa incana* | Hoary alyssum | 93 | 4 | 2 |
| *Rudbeckia triloba ** | Brown-eyed Susan | 0 | 11 | 88 |
| *Solidago rigida *** | Stiff goldenrod | 0 | 45 | 53 |
| *Eryngium yuccifolium ** | Rattlesnake-master | 0 | 10 | 80 |
| *Achillea millefolium* | Common yarrow | 27 | 16 | 42 |
| *Daucus carota ** | Queen Anne’s lace | 0 | 4 | 78 |
| *Silene vulgaris* | Bladder campion | 0 | 65 | 3 |
| *Dalea purpurea ** | purple prairie-clover | 0 | 0 | 65 |

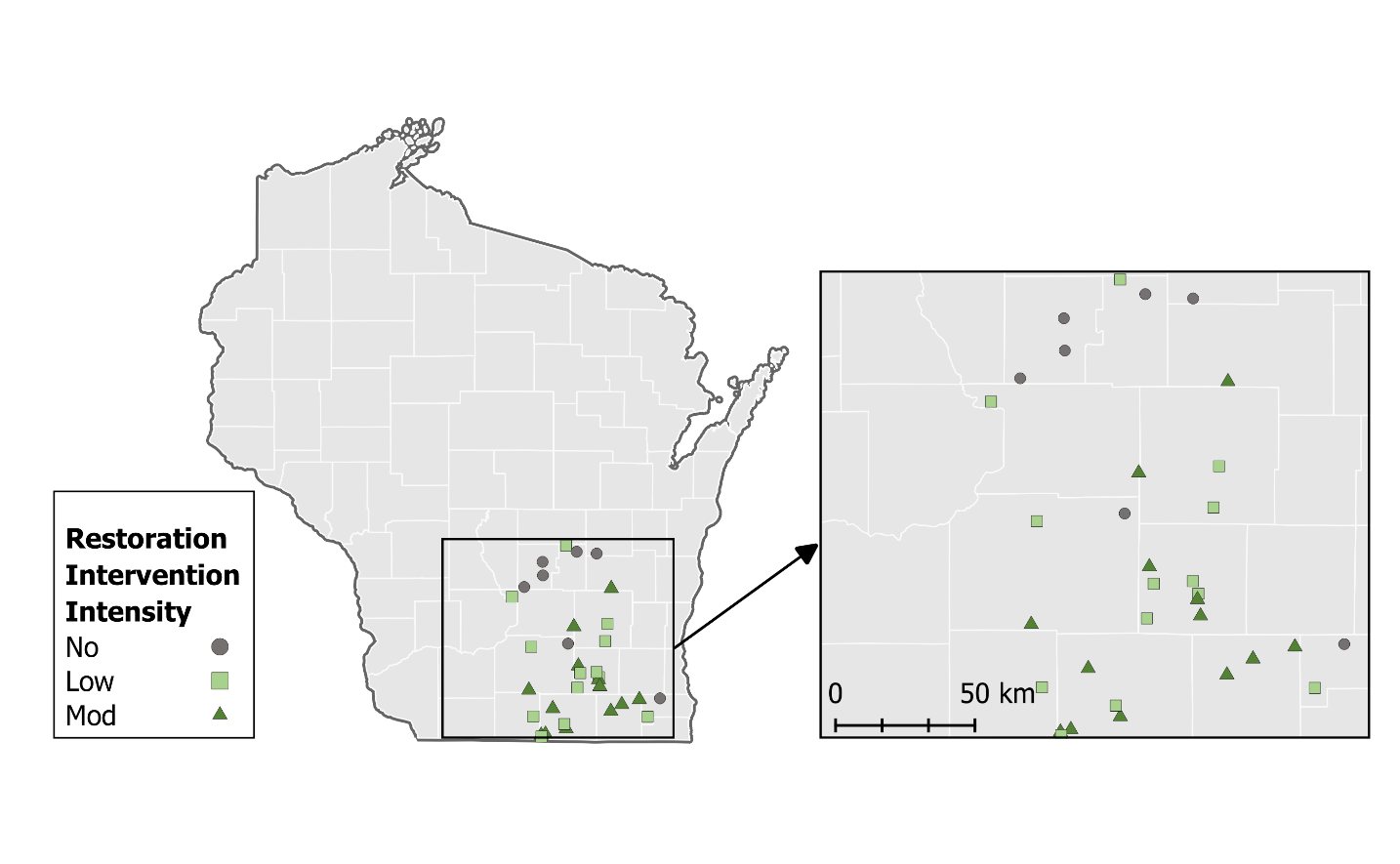

**Figure S1.** Map of the study sites distributed across southern Wisconsin, USA. Shapes and colors of points represent the restoration intervention intensity. Gray circles indicate site with no restoration intervention (*n* = 6), light green squares indicate sites with low intervention intensity (seeded, *n* = 13), and dark green triangles represent moderate intervention intensity (seeded and burned, *n* = 13).

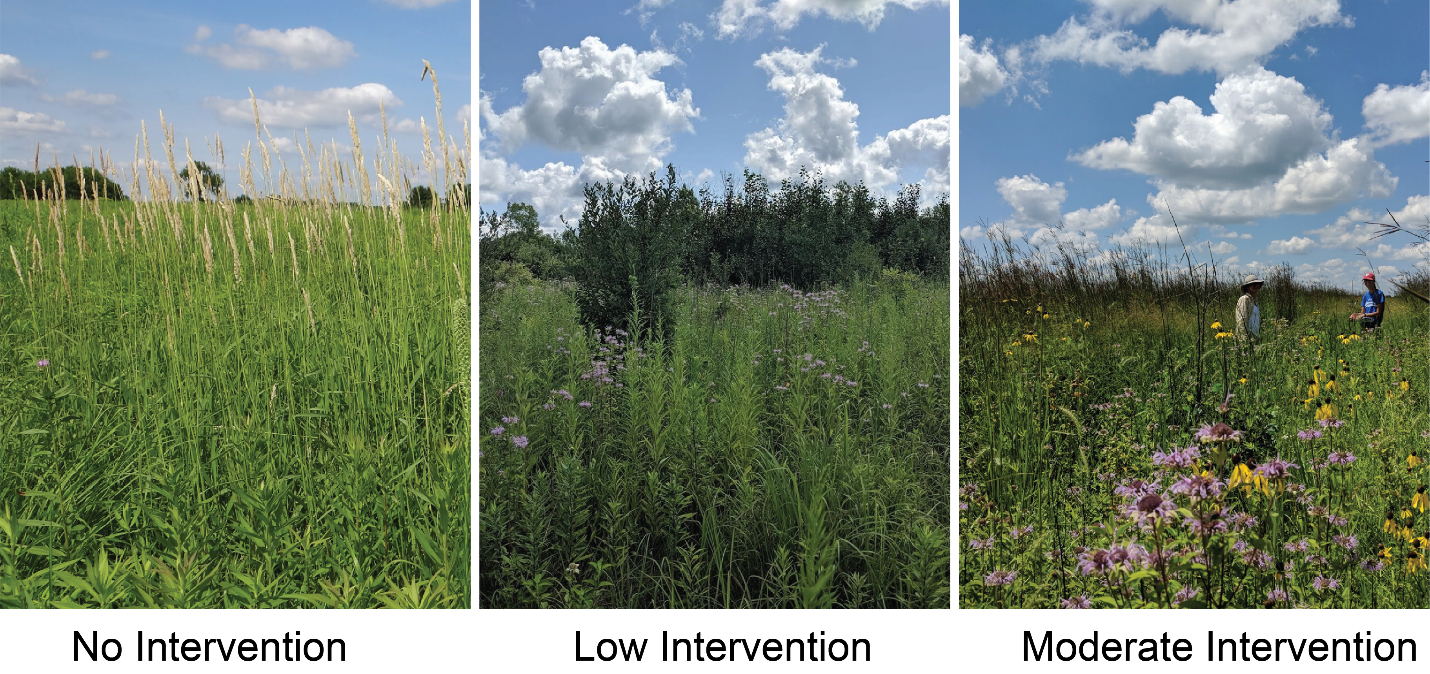
**Figure S2.** Images that represent the typical sites in each category of restoration intervention intensity. Sites without restoration intervention tended to be dominated by cool season grasses (*n* = 6). Sites with low restoration intervention had low floral diversity and tended to have woody shrub encroachment to varying degrees (*n* = 13). Moderate intervention intensity sites typically had abundant and diverse flowering plant communities (*n* = 13). An in-depth analysis of restoration characteristics and plant communities can be found in (McFarlane et al., 2023).

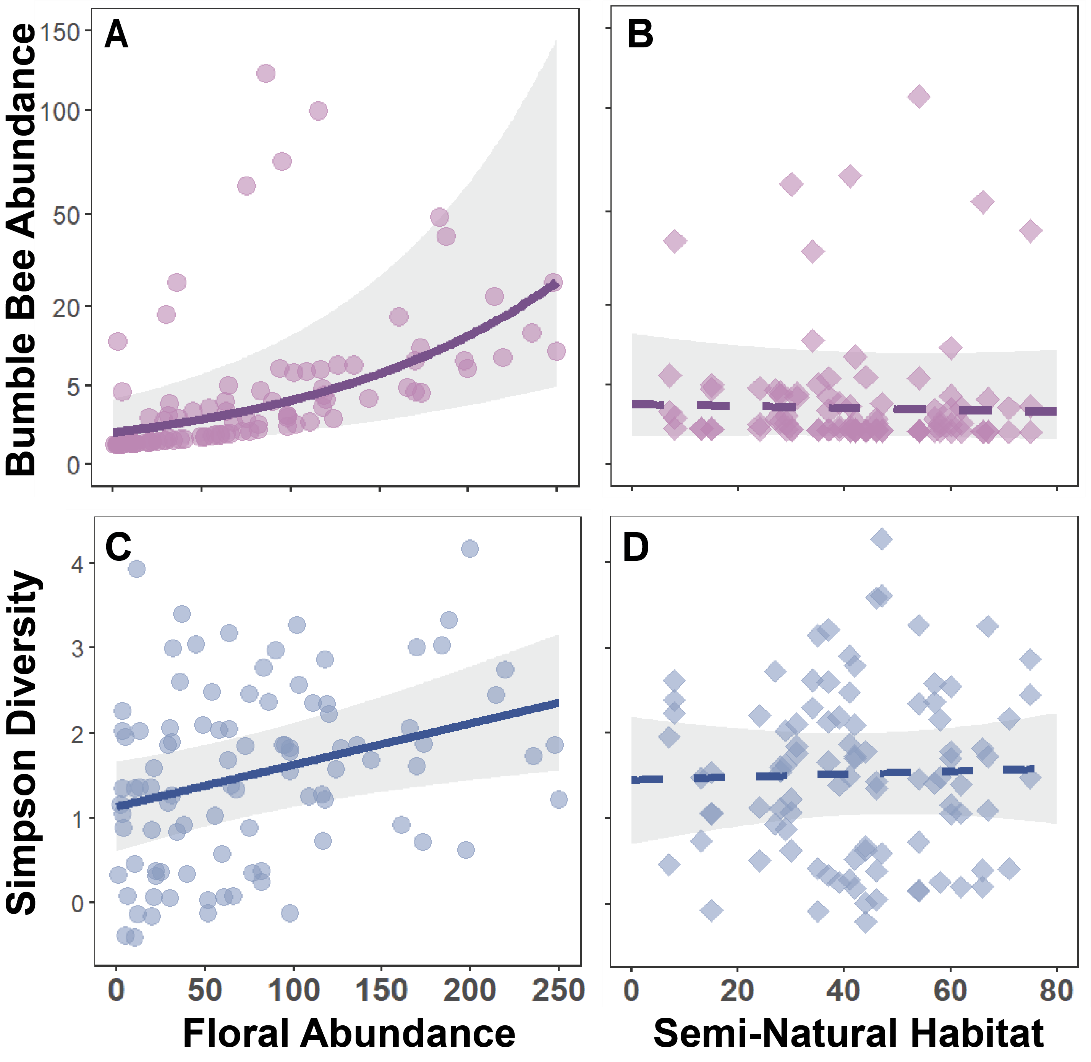

**Figure S3.** Effects of floral abundance and amount of semi-natural habitat in the surrounding 1.5-km on bumble bee abundance and diversity in restored tallgrass prairie. (A) Bumble bee abundance increased with floral abundance (Type II Wald ꭕ² = 26.36, *P* < 0.001), but (B) did not vary with semi-natural habitat (Type II Wald ꭕ² = 0.12, *P* = 0.726). (C) Bumble bee diversity (Inverse Simpson’s Diversity) increased with floral abundance (Type II Wald ꭕ² = 6.42, *P* = 0.040) and (D) did not vary with semi-natural habitat (Type II Wald ꭕ² = 0.07, *P* = 0.786). These plots show partial residuals (±95% CI) after accounting for other fixed and random effects; each point represents total bumble bees or diversity per visit. Lines show GLMM model predictions and shading indicates 95% CI; solid lines are statistically significant slope. Bumble bee abundance is shown on a square-root transformed axis for visual clarity; see methods for the glmm analysis.

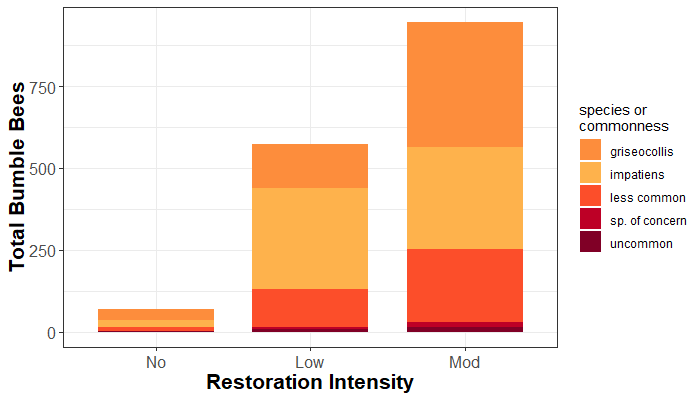

**Figure S4.** A stacked bar graph depicting abundances of bumble bee species compared across restoration intervention intensity. Colors denote the two dominant *Bombus* species and groups of species based on commonness for visual clarity (less common = *B. rufocinctus*, *B. vagans-sandersoni*, *B.* *bimaculatus*; uncommon = *B. citrinus*, *B. borealis*; species of concern = *B. affinis*, *B. fervidus*. Species of concern were only observed at actively restoration (Low and Mod) sites. Community composition significantly differed between restoration intervention intensities (PERMANOVA, *Pseudo-F*_2,36_ = 2.36, *P* = 0.009; Stress = 0.128; Fig. 2). The community compositional differences are likely driven by large changes in abundance (Table S2). Data from both years of study (2018 and 2019) were used.

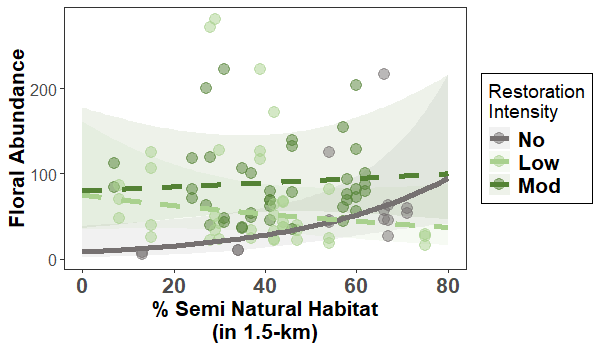

**Figure S5.** Estimated marginal means (± 95% CI) of floral abundance (partial residual effects after accounting for other fixed and random effects in the model). Floral abundance significantly differed with restoration intervention (Type II Wald χ² = 11.52, *P* = 0.003) and the interaction between restoration intervention and the amount of semi-natural habitat in the broad-scale landscape (1.5-km) around study sites (Type II Wald χ² = 6.94, *P* = 0.031).

**Appendix A. Restoration intervention intensity site descriptions.**

Our study sites were restored prairies across Southern Wisconsin, USA, that had management records maintained by the Natural Resources Conservation Service (NRCS). All sites were enrolled in the NRCS Agricultural Conservation Easement Program (ACEP) Wetland Reserve Easements (WRE) and were previously used for row crop agriculture. Management records typically included information on restoration seeding lists, date of restoration, restoration size, and some fire history. We restricted study sites to those within a 161 km (100 mile) radius from Madison, Wisconsin, and that had at minimum 0.8 ha (2 acres) of restored upland. Actively restoration intervention sites (low and moderate intensity) were seeded with dry-mesic, mesic, or wet-mesic plant species depending on site soil and hydrological conditions. Seed mixes were composed of a mix of grasses, forbs, and legumes and species richness ranged from 7 to 54 (mean: 26). The most common grass species included Big bluestem (*Andropogon* *gerardii*), Canada wild rye (*Elymus* *canadensis*), Virginia wild rye (*Elymus* *virginicus*), Indiangrass (*Sorghastrum* *nutans*), and Switchgrass (*Panicum* *virgatum*); less commonly planted species included Little bluestem (*Schizachyrium* *scoparium*), Side-oats grama (*Bouteloua* *curtipendula*), Prairie dropseed (*Sporobolus* *heterolepis*), and Prairie cordgrass (*Spartina* *pectinata*). Most sites were planted only once, though a few were planted twice in different years. Prescribed fires occurred during the dormant season, typically in the spring (Feb – May). Frequency of fire varied across sites within the moderate intervention category; management records were such that burn permits were issued for a site over a range of years so we did not have documentation of burn frequency, intensity, or extent at a given site.
